# A coordinated kinase and phosphatase network regulates Stu2 recruitment to yeast kinetochores

**DOI:** 10.1101/2024.11.01.621564

**Authors:** Michael G. Stewart, Joseph S. Carrier, Jacob A. Zahm, Stephen C. Harrison, Matthew P. Miller

**Affiliations:** Department of Biochemistry, University of Utah School of Medicine, Salt Lake City, United States; Department of Biological Chemistry and Molecular Pharmacology, Harvard Medical School, and Howard Hughes Medical Institute, Boston, United States

## Abstract

Cells coordinate diverse events at anaphase onset, including separase activation, cohesin cleavage, chromosome separation, and spindle reorganization. Regulation of the XMAP215 family member and microtubule polymerase, Stu2, at the metaphase-anaphase transition determines a specific redistribution from kinetochores to spindle microtubules. We show that cells modulate Stu2 kinetochore-microtubule localization by Polo-like kinase1/Cdc5-mediated phosphorylation of T866, near the Stu2 C-terminus, thereby promoting dissociation from the kinetochore Ndc80 complex. Cdk/Cdc28 likely primes Cdc5:Stu2 interaction. Cdc28 activity is also required for Stu2 nuclear import. PP2A^Cdc55^ actively opposes Cdc5 activity on Stu2^T866^ during metaphase. This counter-regulation allows for switchlike redistribution of Stu2^pT866^ at anaphase onset when separase inhibits PP2A^Cdc55^. Blocking Stu2^T866^ phosphorylation disrupts anaphase spindle progression, and we infer that PP2A^Cdc55^ regulates the mitotic spindle by dephosphorylating Stu2 and other MAPs. These data support a model in which increased phosphorylation at anaphase onset results from phosphatase inhibition and point to a larger regulatory network that facilitates rapid cytoskeletal modulation required for anaphase spindle maintenance.

**SUMMARY:** Stu2 displays dynamic localization patterns in the cell cycle, with different kinetochore and microtubule distribution during distinct phases. Phosphorylation near Stu2’s C-terminus reduces its attachment to kinetochores to promote its microtubule activity in anaphase. Cdc5 and PP2A^Cdc55^ play counteracting roles in this pathway to promote proper timing of Stu2 phosphorylation.

## INTRODUCTION

Accurate chromosome segregation during cell division ensures equal partitioning of genetic information into daughter cells. Cells must therefore exert precise temporal control on the complex set of regulatory events that determine proper segregation. At anaphase onset in particular, cells must coordinate separase activation and subsequent removal of cohesin, chromosome movement, spindle reorganization, and preparation for cytokinesis (De Gramont and Cohen-Fix, 2005). Multiple factors, including microtubule associated proteins (MAPs) and motor proteins, are also required during anaphase to maintain the integrity of the microtubule cytoskeleton (Goshima and Vale, 2003; Khmelinskii *et al.*, 2007; Amin, Agarwal and Varma, 2019).

The XMAP215 family member and microtubule polymerase Stu2 is among the proteins that cells must regulate at anaphase onset. Stu2 is a MAP that is essential for proper chromosome segregation, performing multiple important roles in the cell. Stu2 binds to the plus ends of microtubules to regulate microtubule dynamics (Kosco *et al.*, 2001; van Breugel, Drechsel and Hyman, 2003; Al-Bassam *et al.*, 2006; Ayaz *et al.*, 2012, 2014), but it also must localize to kinetochores through an interaction with the Ndc80 complex (Ndc80c) to carry out functions essential for chromosome biorientation (Hsu and Toda, 2011; Tang *et al.*, 2013; Miller, Asbury and Biggins, 2016; Vasileva *et al.*, 2017; Miller *et al.*, 2019; Herman, Miller and Biggins, 2020; Zahm *et al.*, 2021). It also has important functions at microtubule organizing centers to support microtubule nucleation (Wang and Huffaker, 1997; Gunzelmann *et al.*, 2018) and at the spindle midzone to promote anaphase spindle elongation (Severin *et al.*, 2001). This diversity of function requires dramatic relocalization of Stu2 during the cell cycle. Fluorescently labeled Stu2 shifts between kinetochores and microtubules as well as into and out of the nucleus at different cell cycle stages, (Usui *et al.*, 2003; Ma *et al.*, 2007; Van Der Vaart *et al.*, 2017), and photobleaching studies show changes in Stu2 dynamics at kinetochores from metaphase to anaphase (Aravamudhan *et al.*, 2014). Understanding how cells alter Stu2’s activity and localization in the cell cycle is important for uncovering how it facilitates proper spindle maintenance and chromosome segregation, and how aneuploidy might result when these processes go awry.

Strict coordination with the cell cycle implicates phosphoregulation by cell-cycle kinases. We show here that phosphorylation by cyclin dependent kinase (Cdk/Cdc28 in yeast) of a conserved serine in Stu2’s nuclear localization signal (NLS) promotes nuclear import. Once in the nucleus, Stu2 associates predominantly with kinetochores through its interaction with the Ndc80c. At anaphase onset, phosphorylation of a conserved threonine near the C-terminus of Stu2 (Stu2^T866^) reduces the amount of Stu2 bound to Ndc80c. We show that Stu2^T866^ phosphorylation depends on Polo-like kinase 1 (Plk1/Cdc5 in yeast) and that phosphorylation of a conserved serine (Stu2^S603^) in the basic-linker region of Stu2 primes it for Cdc5 interaction. Phosphorylation of Stu2^S603^ is the same modification that promotes nuclear import, showing multiple functions for this region of Stu2’s basic linker. The phosphatase PP2A^Cdc55^ opposes Cdc5 modification of Stu2^T866^ during metaphase and sets up proper timing of Stu2^T866^ modification at anaphase onset. Phosphorylation of Stu2^T866^ corresponds with relocalization of a pool Stu2 from kinetochores to interpolar spindle microtubules. Blocking this modification leads to defects in anaphase spindle progression. Furthermore, disrupting PP2A^Cdc55^ activity leads to spindle defects in metaphase and anaphase, indicating this phosphatase broadly regulates the microtubule cytoskeleton during mitosis. These findings illustrate an interconnected and likely conserved network of kinases and phosphatases that regulate Stu2 activity, along with many other factors, to ensure precise timing of the numerous events that unfold in rapid succession at anaphase onset.

## RESULTS

### Stu2:Ndc80c association changes in the cell cycle

To assess changes in the association of Stu2 and Ndc80c across the cell cycle, we measured signal from fluorescently labeled Stu2 and Ndc80, in yeast strains grown synchronously from a G1 arrest-release. For this and other in vivo assays, we made use of an Auxin Inducible Degron (AID), in which the endogenous *STU2* locus is fused with IAA7 (hereafter *stu2-AID*) and degraded upon treatment with auxin, uncovering phenotypes of ectopic mutants or fusion proteins (Nishimura *et al.*, 2009). We released G1-arrested *stu2-AID NDC80-mKate* cells harboring ectopic *STU2-GFP* and monitored both total cellular Stu2-GFP signal as well as the kinetochore-localized Stu2-GFP throughout the cell cycle (Fig. 1A). To quantify the kinetochore-localized Stu2-GFP signal, we calculated the ratio of the intensities of Stu2-GFP and Ndc80-mKate in kinetochore puncta, as in previous studies (Aravamudhan *et al.*, 2014). Because Ndc80c is Stu2’s kinetochore receptor, changes in the Stu2/Ndc80c ratios will reflect dynamic regulation. Total cell Stu2-GFP levels rise steadily through G1/S phases and reach a peak in mitosis. This trend is consistent with previous reports that total Stu2 expression increases as cells progress through the cell cycle (Guo *et al.*, 2006; Santos, Wernersson and Jensen, 2015). The kinetochore-associated Stu2-GFP pool also increases through G1/S phases, mirroring total Stu2-GFP, but as the cells enter early mitosis, which marks the peak in the level of total cell Stu2-GFP, kinetochore-associated Stu2 decreases substantially (time point 75 minutes, Fig. 1A, Fig. S1A). Both total cell and kinetochore Stu2-GFP levels then fall and reach stable signal in the new G1 phase. These data suggest that during M phase, specific mechanisms cause Stu2 to dissociate from Ndc80c, despite the continuing rise in total Stu2.

**Figure 1.**
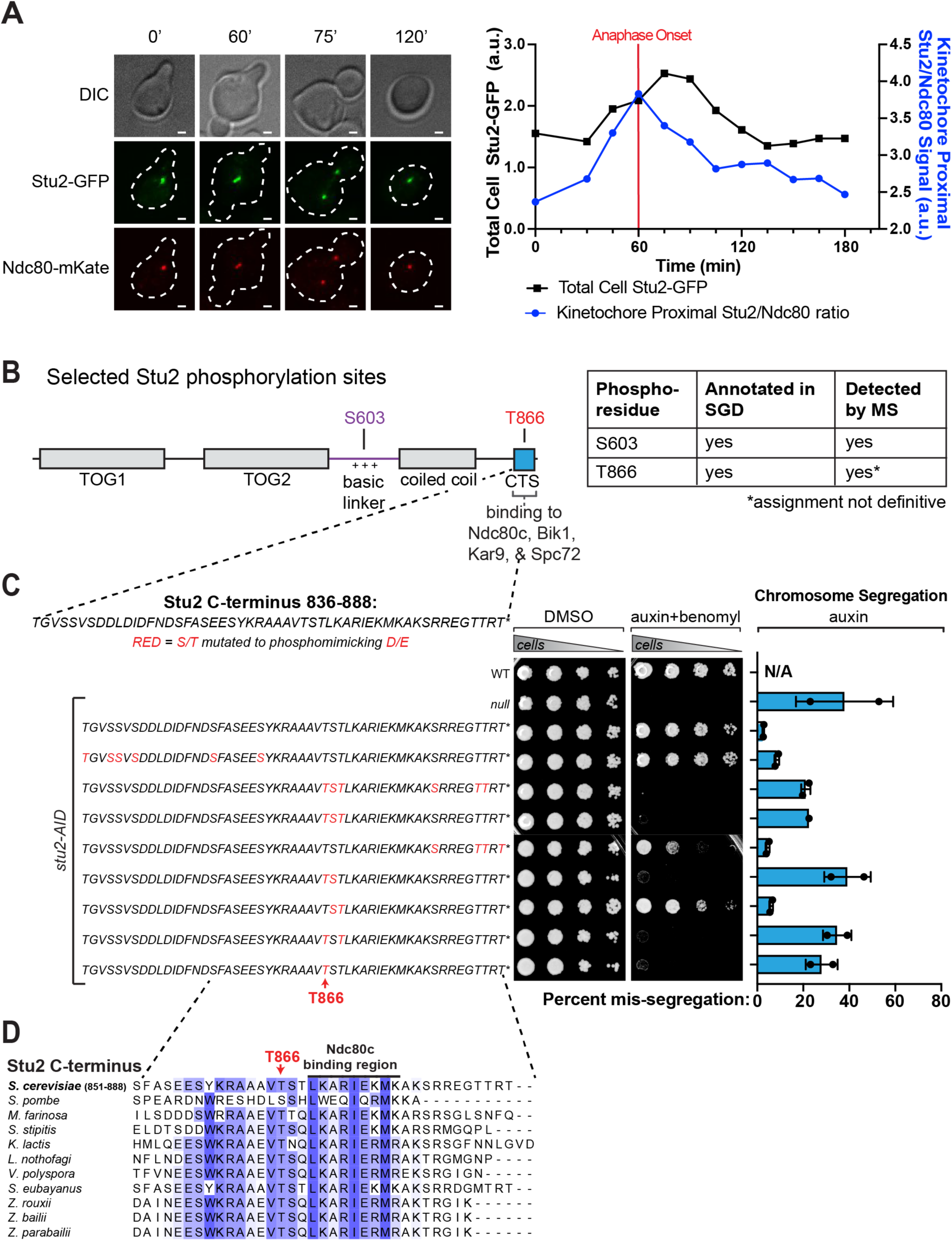
Evidence for regulation of Stu2-Ndc80c assembly in the cell cycle and identification of phosphorylated Stu2 T866. A. Exponentially growing *stu2-AID* cells expressing *STU2-GFP and NDC80-mKate* (M3774), were released from a G1 arrest into auxin-containing media. Samples were taken every 15 minutes, fixed, and imaged. LEFT: Representative images of cells fixed at indicated time points. Scale bar is 1 µm. RIGHT: Total GFP signal from individual cells was quantified for each time point (black line). Kinetochore Stu2-GFP signal and Ndc80-mKate signal were measured for individual puncta at each time point. Ratio of Stu2/Ndc80c signal plotted over the time course (blue line). All measurements for each time point (Total Cell-Stu2-GFP, Kinetochore Stu2/Ndc80 ratio) are an average from 2 replicate time course experiments. Total number of cells for each measurement range from 184-236. Data points are mean. Error bars are S.E.M, but error values are too small to easily visualize. B. Illustrated residues on the domain-map of Stu2 indicate phosphorylated Serine and Threonine residues identified by mass-spectrometry (for full list of identified residues, see Fig. S1B). Exponentially growing *STU2-3FLAG* (M498) cultures were harvested, lysed to produce protein sample, subjected to α-Flag IP, and analyzed by mass-spectrometry. Table shows which residues were also previously annotated in Saccharomyces Genome Database (SGD). C. LEFT - Wild-type (M3), *stu2-AID* (no covering allele, M619), and *stu2-AID* cells expressing various *STU2-3HA* alleles from an ectopic locus (*STU2^WT^*, M2898; *stu2^T836E S839D S840D S842D S852D S855D S858D^*, M3352; *stu2*^*T866E S867D T868E S880D T885E T886E*^, M3353; *stu2*^*T866E S867D T868E*^, M3354; *stu2*^*S880D T885E T886E T888E*^, M3355; *stu2*^*T866E S867D*^, M3356; *stu2*^*S867D T868E*^, M3357; *stu2^T866E T868E^*, M3358; *stu2^T866E^*, M2829) were serially diluted (five-fold) and spotted on plates containing DMSO (control) or 500 μM auxin + 5 μg/mL benomyl. Residues shown in red indicate S/T amino acids mutated to D/E. RIGHT – Exponentially growing wild Type cells (M3) and *stu2-AID* cells harboring *(CEN3.L.YA5.1)MATα* on a mini-chromosome as well as *MFA1-3xGFP,* with or without *STU2-3HA* covering alleles, (no covering allele, M3276; *STU2^WT^,* M3451; *stu2^T836E S839D S840D S842D S852D S855D S858D^,* M3592; *stu2*^*T866E S867D T868E S880D T885E T886E*^, M3593; *stu2*^*T866E S867D T868E*^, M3594; *stu2*^*S880D T885E T886E T888E*^, M3595; *stu2*^*T866E S867D*^, M3596; *stu2*^*S867D T868E*^, M3597; *stu2*^*T866E T868E*^, M3598; *stu2*^*T866E*^, M3452) were diluted into non-selective media containing 500 μM auxin. Cells were cultured for 16 hours, fixed, and then analyzed by flow cytometry to determine the percentage of cells mis-segregating the mini-chromosome. Bars are mean of 2 biological replicates. Error bars are S.E.M. D. Multiple sequence alignment of the Stu2 C-terminus and C-termini from Stu2 fungal homologs. Hydrophobic residues important for Ndc80c binding (Zahm *et al.*, 2021) are indicated under the labeled black line.

### Phosphorylation of T866 near the Stu2 C-terminus

Reversible phosphorylation regulates the function of many kinetochore proteins, especially by modulating their association with larger complexes (Ciferri *et al.*, 2008; Sundin, Guimaraes and DeLuca, 2011; London *et al.*, 2012; Sarangapani *et al.*, 2013; Zaytsev *et al.*, 2015; Jenni and Harrison, 2018; Gutierrez *et al.*, 2020; Dudziak *et al.*, 2021). We have shown previously that the C-terminal segment (CTS) of Stu2 binds the tetramerization junction of the Ndc80 complex (Zahm *et al.*, 2021). The CTS contains several serine and threonine residues, and we hypothesized that phosphorylation of the CTS could affect its association with Ndc80c. To determine if any residues in the CTS were phosphorylated, we conducted phosphoproteomic analysis of Stu2 purified from yeast. We detected many phosphorylated Stu2 peptides, including one containing CTS residues T866, S867, and T868 (Fig. 1B, Fig. S1B). Phosphorylation at T866 was previously reported in a proteome-wide study from yeast (Lanz *et al.*, 2021). To determine which of the CTS residues is phosphorylated in vivo, we turned to a phenotypic screen.

Our previous work showed that mutations which perturb the binding of Stu2’s C-terminus with Ndc80c result in cellular growth and chromosome segregation defects (Zahm *et al.*, 2021). We made phosphomimetic substitutions throughout the Stu2 CTS by replacing serines and threonines with aspartates and glutamates, respectively. We then examined the cell growth phenotype of these mutations expressed ectopically in *stu2-AID* cells (Fig. 1C). We generated combinations of mutations and found that cells harboring the phosphomimetic T866E mutation were defective in growth on auxin, especially in the presence of the microtubule poison benomyl (Fig. 1C, Fig. S1C). We also examined chromosome segregation phenotypes of the same panel of phosphomimetic mutants using a quantitative Chromosome Transmission Fidelity assay (Zhu *et al.*, 2015) (Fig. 1C, Fig. S1D). Mutant constructs harboring T866E showed chromosome segregation defects that aligned with cell growth defects. Highlighting the importance of T866 in vivo, multiple sequence alignments also show that T866 is conserved across fungal species (Fig. 1D). These results, along with the MS data described above, support the idea that Stu2’s CTS is phosphorylated at T866 in yeast cells.

### Phosphorylation of Stu2^T866^ reduces its association with Ndc80c

T866 is adjacent to the Ndc80c binding region of Stu2 (Fig. 1D). Inspection of the structure shows that it lies above a hydrophobic patch that mediates interaction with the CTS (Fig 2A-B) (Zahm *et al.*, 2021). Modeling a phosphothreonine at T866 (pT866) appears to place a highly charged side chain unfavorably close to this hydrophobic surface (Fig. 2B). Moreover, neighboring surfaces of Ndc80c have a net negative charge, likely to further destabilize association of Stu2 bearing a phosphate at T866 (Fig. S2A). As shown in Fig. 2C-D, fluorescence polarization binding experiments, in which we measured displacement of an Oregon Green labeled Stu2 CTS peptide from Ndc80c^Dwarf^ by an unlabeled CTS peptide, with either unmodified T866 or pT866, showed about a 6-fold lower affinity of the phosphorylated peptide (unmodified 34 ± 7.2 μM vs pT866 200 ± 70 μM). A control scrambled C-terminus peptide sequence showed no detectable binding to Ndc80^Dwarf^ (Fig. 2D). These data provide direct evidence that pT866 lowers affinity of Stu2 for Ndc80c.

**Figure 2.**
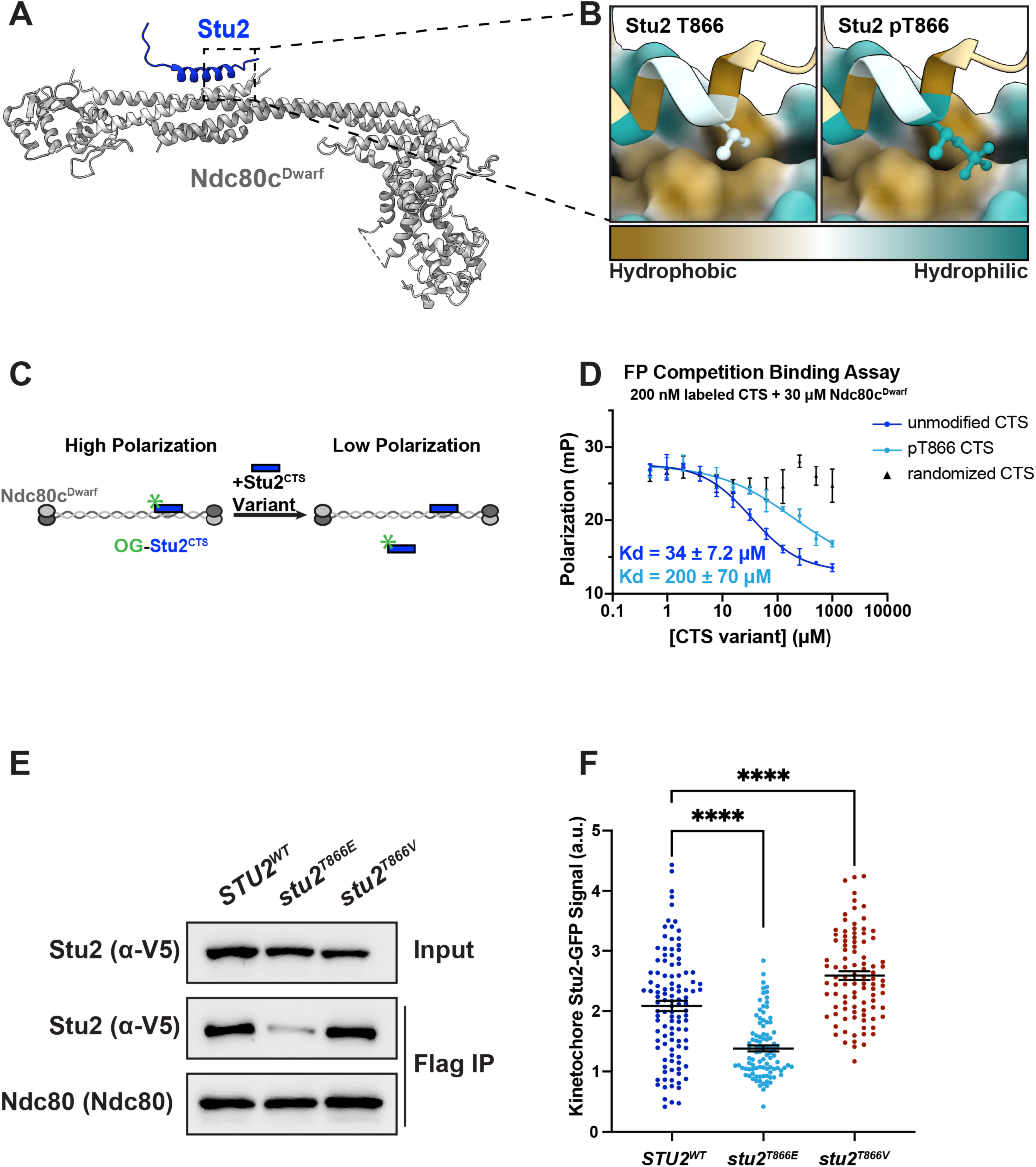
Phosphorylated T866 decreases binding of Stu2 to Ndc80c. A. The crystal structure of Stu2 C-terminus bound to Ndc80^Dwarf^ (PDB 7KDF); see (Zahm *et al.*, 2021). Ndc80c^Dwarf^ in grey, Stu2 C-terminus in blue. B. Zoom in of crystal structure showing hydrophobicity of Stu2 and Ndc80c amino acids rendered using ChimeraX on the Kyte-Doolittle scale (kdHydrophobicity). Ndc80c proteins in surface representation; Stu2 in ribbon representation. LEFT – unmodified Stu2^T866^ as in PDB 7KDF. RIGHT – Modeling of Stu2^pT866^. C. Schematic representation of fluorescence polarization competition assay. Oregon Green-labeled Stu2 CTS peptide pre-bound to Ndc80^Dwarf^ then treated with unlabeled Stu2 peptide with or without pT866 and fluorescence polarization was measured. D. A Stu2 CTS peptide (855-888), a Stu2 CTS peptide containing a phosphothreonine at position 866, and a peptide with a randomized CTS sequence were subjected to serial 2-fold dilutions, and each dilution was mixed 1:1 with a solution containing 30 µM Ndc80^Dwarf^, and 200 nM Oregon green-labeled CTS peptide, also in resuspension buffer. Fluorescence polarization was measured in triplicate in 96-well plates. Error bars are S.D. Kd calculated with nonlinear fitting in Prism. E. Exponentially growing *stu2-AID* cultures expressing an ectopic copy of *STU2* (*STU2^WT^*, M622; *stu2^T866E^*, M1448; *stu2^T866V^*, M4398) as well as *DSN1-6His-3Flag* at the genomic locus were treated with auxin 30 min prior to harvesting. Kinetochore particles were purified from lysates by anti-Flag immunoprecipitation (IP) and analyzed by immunoblotting. F. Exponentially growing *stu2-AID, pMET-CDC20* cultures with an ectopically expressed *STU2-GFP* allele (*STU2^WT^-GFP*, M2599; *stu2^T866E^-GFP*, M2600; *stu2^T866V^-GFP*, M4447) that also contained *SPC110-mCherry* (spindle pole) were shifted to auxin-containing media to degrade Stu2-AID and supplemented with methionine to arrest cells in metaphase by depleting Cdc20. Cells were fixed and imaged to determine Kinetochore-Proximal Stu2-GFP. Bars represent mean of n=96-111 individual measurements. Error bars are S.E.M. p-values from two-tailed unpaired t-tests (*STU2^WT^* vs. *stu2^T866E^* p<0.0001; *STU2^WT^* vs. *stu2^T866V^* p=0.0162.)

To assess the effect of T866 mutations on Ndc80c binding in yeast, we continued to use *stu2^T866E^* to mimic constitutively phosphorylated T866 and also generated a *stu2^T866V^* allele to prevent T866 phosphorylation. A valine mutation preserves an interaction between the gamma carbon of Stu2 T866 and Spc24 F14 seen in the crystal structure (Fig. S2B); a smaller residue would leave an unfavorable hole, consistent with the phenotype of a *stu2^T866A^* mutant (Fig. S2B-E). We purified kinetochores from asynchronously growing *stu2-AID* yeast harboring *STU2^WT^-V5, stu2^T866E^-V5,* and *stu2^T866V^-V5* by immunoprecipitation of the kinetochore component Dsn1 (Akiyoshi *et al.*, 2010). Consistent with the fluorescence polarization results, we observed less kinetochore associated Stu2^T866E^ than Stu2^WT^ (Fig. 2E). We also examined the association of Stu2-GFP with kinetochores in metaphase-arrested cells. Metaphase cells have bilobed clusters of Stu2-GFP, closely corresponding to kinetochore localization (Fig. 1A). These clusters have diminished intensity in *stu2* mutants that cannot bind Ndc80c (Zahm *et al.*, 2021). Metaphase-arrested cells expressing *stu2^T866E^-GFP* had lower levels of kinetochore Stu2 than *STU2^WT^-GFP* (Fig. 2F) and *stu2^T866V^-GFP* had higher levels (Fig. 2F). These results are consistent with the conclusion that Stu2^T866^ phosphorylation reduces Stu2 association with Ndc80c.

### Cdc5 phosphorylates Stu2^T866^

Which kinase phosphorylates Stu2 T866? Standard bioinformatics tools (Wang *et al.*, 2020) produced no strongly matched kinase consensus motif, but the apparent timing of this modification implied that we should search for a mitotic kinase. We therefore devised a fluorescence microscopy assay to assess Stu2:Ndc80c binding after local activation of a panel of mitotic kinases at the kinetochore. In *STU2-GFP* cells, we made fusions of the mitotic kinases to the TOR subunit FRB and also expressed *NUF2-FKBP*, so that we could use rapamycin to recruit each kinase to Ndc80c (Fig. 3A) (Haruki, Nishikawa and Laemmli, 2008). The C-terminus of Nuf2 is very close to the endogenous Stu2 binding site on Ndc80c, so recruiting the correct kinase to Nuf2 should result in phosphorylation of Stu2 and reduction of Stu2-GFP signal at the kinetochore, as we observed in the phosphomimetic *stu2^T866E^* cells (Fig. 2F). To remove any confounding effect of cell cycle regulation of these kinases, we performed these assays in cells arrested in G1 or metaphase. Recruitment of Bub1-FRB, Mps1-FRB, and Ipl1-FRB to kinetochores resulted in no changes in Stu2-GFP levels in either G1 or mitotically arrested cells (Fig. S3A-B). Recruitment of Cdc5-FRB to kinetochores resulted in a significantly lower Stu2-GFP signal that matched the magnitude of signal reduction we observed in *stu2^T866E^* cells (Fig. 3B compare Fig. 2F). Cdc5-FRB recruitment to kinetochores decreased Stu2-GFP signal in both G1 and mitotically arrested cells (Fig. 3B and Fig. S3C) and depended on the presence of Nuf2-FKBP and dosage of rapamycin (Fig. S3D-E). Cells harboring *stu2^T866V^-GFP* showed no reduction of kinetochore Stu2-GFP signal upon tethering of Cdc5-FRB to kinetochores, consistent with T866 as the critical Cdc5 target (Fig. 3B, Fig. S3C-D). Additional evidence for the role of Cdc5 came from knockdown experiments using the *cdc5-1* temperature sensitive allele. Mitotically arrested cells were shifted to non-permissive 37°C for 1 hour. *cdc5-1* cells had higher Stu2-GFP signal at the kinetochore than did *CDC5* control cells, indicating that the removal of Stu2 from the kinetochore depended on Cdc5 activity (Fig. 3C). Finally, we examined the cellular effect of ectopic Cdc5 recruitment to kinetochores. Tethering Cdc5-FRB to Nuf2-FKBP resulted in a severe growth defect that could be rescued by expression of *stu2^T866V^* (Fig. 3D, Fig. S3F). These results all support our conclusion that Cdc5 phosphorylates Stu2 at T866.

**Figure 3.**
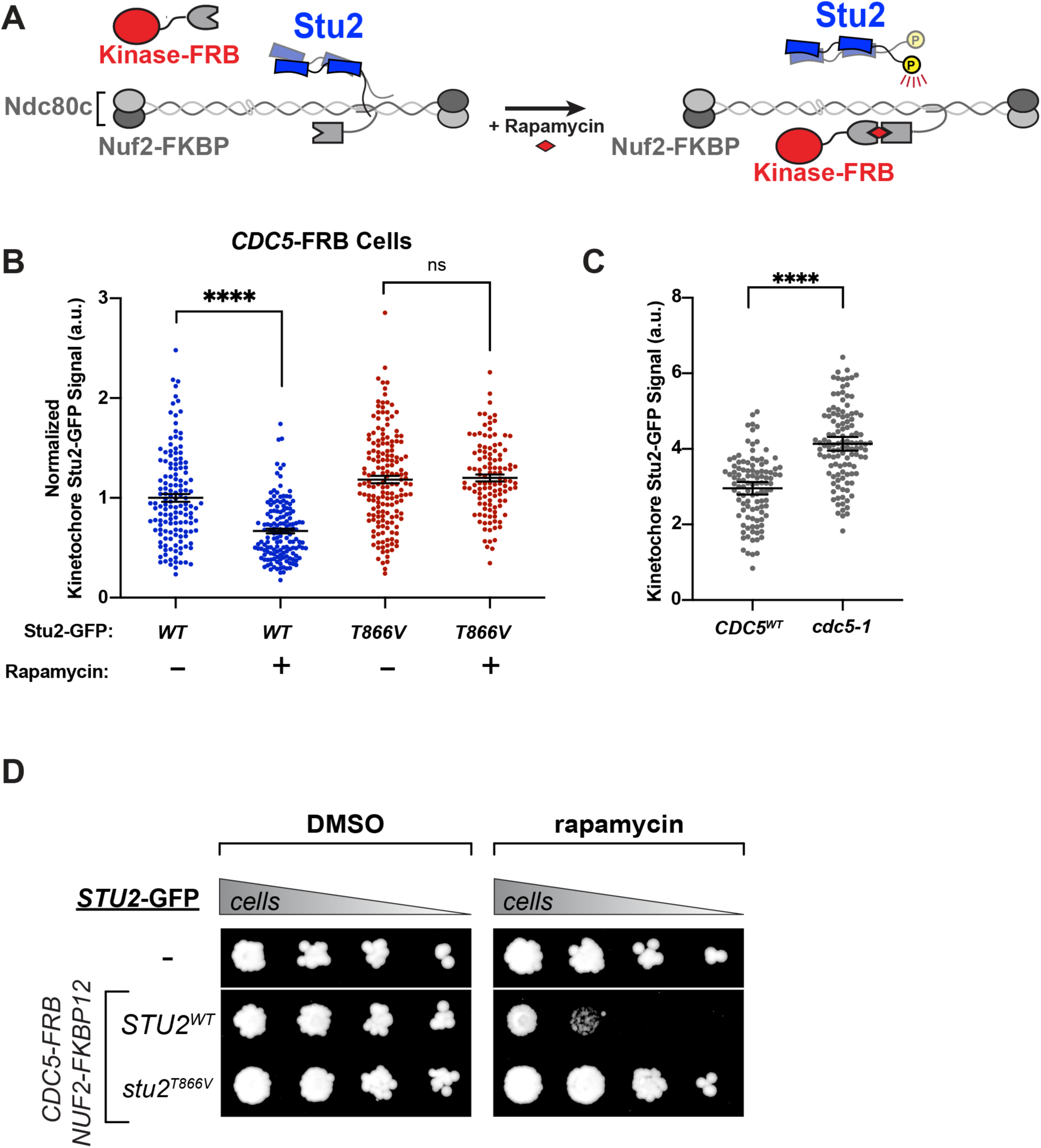
Cdc5 phosphorylates Stu2^T866^. A. Schematic illustrating activation of kinases at the kinetochore by chemical-genetic tethering. Ndc80 and Spc24 shown in light grey, Nuf2 and Spc25 are shown in dark grey. In the presence of rapamycin, the kinase-FRB fusion is induced to bind Nuf2-FKBP12, localizing the kinase to the kinetochore. If the kinase is able to phosphorylate Stu2^T866^, we postulated that Stu2 will unbind Ndc80c and delocalize from the kinetochore. B. Exponentially growing *stu2-AID* cells harboring *TOR1-1, fpr1Δ, NUF2-FKBP12, CDC5-FRB,* and an ectopic *STU2-GFP* variant (*STU2^WT^-GFP,* M4968; *stu2^T866V^-GFP,* M4969) were arrested in alpha factor for 3 hours. Cells received either 500 µM auxin + 200 ng/mL rapamycin or 500 µM auxin + DMSO for 30 minutes prior to being fixed and imaged. Stu2-GFP puncta intensity was quantified using ImageJ. Bars are average of n=114-166 individual measurements. Error Bars are S.E.M. p-value from an unpaired t-test (*STU2^WT^* + DMSO vs *STU2^WT^* + RAP, p < 0.0001; *stu2^T866V^* + DMSO vs *stu2^T866V^* + RAP, p=0.763). C. Exponentially growing cells harboring endogenous *STU2-GFP* and *pMET-CDC20,* with or without *cdc5-1* temperature sensitive allele (*CDC5^WT^,* M2827; *cdc5-1*, M3138), were cultured in methionine-containing media to arrest cells in metaphase for 2 hours. Cells were then transferred to 37°C to inhibit *cdc5-1* for 1 hour. Cells were fixed and imaged to determine kinetochore-proximal Stu2-GFP. Bars are average of n=106-116 individual measurements. Error bars are S.E.M. p-value from two-tailed unpaired t-tests (*CDC5^WT^* vs *cdc5-1*, p<0.0001). D. Cells harboring *TOR1-1* & *fpr1Δ (*M1375), and *TOR1-1 fpr1Δ NUF2-FKBP12 CDC5-FRB stu2-AID* cells with either *STU2^WT^-GFP* (M4968) or *stu2^T866V^-GFP* (M4969) were serially diluted and spotted on plates containing DMSO, 500 μM auxin, 0.05 μg/mL rapamycin, or 500 μM auxin + 0.05 μg/mL rapamycin.

### Stu2 basic linker interacts with Cdc5 polo-box domain

Cells have diverse mechanisms to ensure proper targeting of many Cdc5 substrates. One such mechanism is “priming” phosphorylation, in which a substrate must first be phosphorylated at a distal site before Cdc5 can act upon it (Elia *et al.*, 2003; Örd *et al.*, 2020; Singh *et al.*, 2021). For such substrates, the priming site of the substrate frequently consists of a Cdk/Cdc28 phosphorylation site that when phosphorylated interacts with the polo box domains of Cdc5 with high affinity (Fig. S4A). Recent work suggests that in addition to binding to phosphorylated serines/threonines, the polo-box domains harbor an additional binding interface on the face distal to the phosphopeptide binding surface, which interacts with hydrophobic residues in a substrate (Sharma *et al.*, 2019; Almawi *et al.*, 2020). It is not known whether substrates interact with both the phosphopeptide binding surface and hydrophobic surface at the same time. Stu2 harbors a consensus Cdc28 phosphorylation site in its basic linker region, comprising residues 600-607 (Fig. S4A). Both we and others have detected phosphorylation of the consensus Cdc28 site (serine 603) by mass spectrometry (Fig. 1B; see also (Aoki *et al.*, 2006; Humphrey, Felzer-Kim and Joglekar, 2018)). Does Stu2’s basic linker region mediate an interaction with the Cdc5 polo-box domains? We used AlphaFold 3 to generate a predicted structure of the interaction of the Stu2 basic linker region (Stu2^560-657(pS603)^) with the Cdc5 polo-box domains. The predicted structure shows two putative Stu2:Cdc5 interacting regions (Fig. 4A-B). One region, Stu2^592-607^, binds to the phophopeptide binding domain through pS603 and shows strong similarity with Spc72, another phosphorylated Cdc5 substrate previously determined by X-ray crystallography (Fig. S4B) (Almawi *et al.*, 2020). This prediction suggests that pS603 contacts Cdc5 H641 and K643, which are critical for phosphorylated substrate binding (Fig. 4B). The second predicted interaction region is between another conserved basic linker patch (Stu2^634-639^) and the Cdc5 polo-box domain at the hydrophobic binding site (Fig. 4B). This binding mode is similar to another Cdc5 interactor (Dbf4) that binds the hydrophobic site (Fig. S4B) (Almawi *et al.*, 2020). AlphaFold makes these predictions with high confidence based on plDDT and pTM metrics, and a non-phosphorylated Stu2^592-609^ is not predicted to bind Cdc5 polo-box domains (Fig. S4C-D). Both the phospho-binding motif and the hydrophobic binding motif of the Stu2 basic linker are conserved in fungi (Fig. 4A).

**Figure 4.**
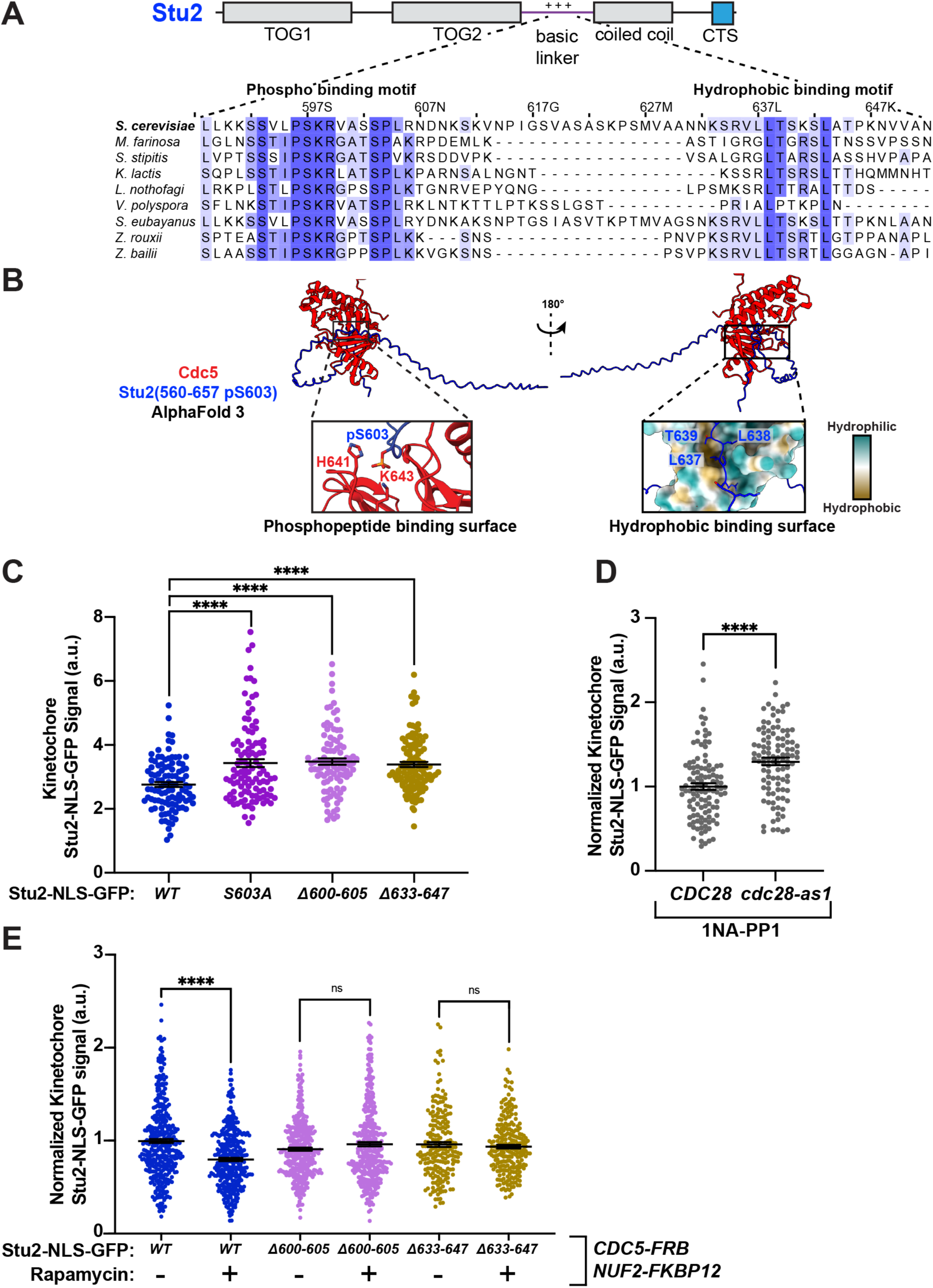
Stu2 basic linker regions mediate interaction between Stu2 and Cdc5. A. Schematic of Stu2 domain structure as well as multiple sequence alignment of fungal species showing conservation of two patches of Stu2 basic linker region. 592-607 is predicted to interact with Cdc5 phoshpopeptide binding surface, while 633-648 is predicted to interact with Cdc5 hydrophobic binding surface. B. AlphaFold 3 prediction of Stu2 basic linker with pS603 (Stu2^560-657 pS603^) bound to Cdc5 polo box domain. Left shows pS603 binding to phosphopeptide binding residues on Cdc5. Right shows hydrophobic residues interacting with a hydrophobic patch on Cdc5 C. Exponentially growing *stu2-AID cdc20-AID* cultures with an ectopically expressed *STU2-NLS^SV40^-GFP* allele (*STU2^WT^-NLS-GFP*, M5145; *stu2^S603A^-NLS-GFP*, M5147; *stu2^Δ600-605^-NLS-GFP, M5149*; *stu2^Δ633-647^-NLS-GFP*, M5980) that also contained *SPC110-mCherry* (spindle pole) were cultured in auxin-containing media to arrest cells in metaphase. Cells were fixed and imaged to determine Kinetochore-Proximal Stu2-GFP. Bars represent the average of n=104-112 individual measurements. Error bars are S.E.M. p-values from two-tailed unpaired t-tests (*STU2^WT^* vs *stu2^S603A^*, p<0.0001; *STU2^WT^* vs *stu2^Δ600-605^,* p<0.0001; *STU2^WT^* vs *stu2^Δ633-647^-NLS-GFP*, p<0.0001). D. Exponentially growing *stu2-AID* cells with ectopically expressed *STU2-NLS-GFP* and either *CDC28^WT^* (M5554) or *cdc28-as1* (M5555) were arrested in mitosis by nocodazole treatment, then treated with 500 µM auxin and 2.5 µM 1NA-PP1 for 30 minutes prior to harvesting. Cells were fixed and imaged to determine Kinetochore-Proximal Stu2-GFP. Bars represent the average of n=103-112 individual measurements. Error bars are S.E.M. p-values from two-tailed unpaired t-tests (*CDC28^WT^* vs *cdc28-as1*, p<0.0001). E. Exponentially growing *stu2-AID* cells harboring *TOR1-1 fpr1Δ NUF2-FKBP12 CDC5-FRB* and an ectopic *STU2-NLS^SV40^-GFP* variant (*STU2^WT^-NLS-GFP*, M5150; *stu2^Δ600-605^-NLS-GFP*, M5154; *stu2^Δ633-647^-NLS-GFP*, M5989) were arrested in alpha factor for 3 hours. Cells received either 500 µM auxin + 200 ng/mL rapamycin or 500 µM auxin + DMSO for 30 minutes prior to fixation and imaging. Bars represent the average of 3 biological replicates normalized to *STU2^WT^* and n=205-374 individual measurements. Outliers were removed from data by ROUT analysis with Q=1%. Error bars are S.E.M. p-value from an unpaired t-test (*STU2^WT^* + DMSO vs *STU2^WT^+* RAP, p<0.0001; *stu2^Δ600-605^* + DMSO vs *stu2^Δ600-605^* + RAP, p=0.0508; *stu2^Δ633-647^* + DMSO vs *stu2^Δ633-647^* + RAP, p=0.9586).

If Stu2^T866^ is phosphorylated by Cdc5 to regulate its interaction with Ndc80c, several predictions emerge. First, mutating either of the putative polo-box binding motifs in Stu2’s basic linker would prevent interaction with Cdc5’s polo-box domain. As a result, Stu2 would not be phosphorylated at T866, leading to higher kinetochore localization, similar to Stu2^T866V^. Second, removal of Stu2 from kinetochores would require Cdk-dependent priming. To test the first prediction, we made mutants to both conserved regions of Stu2’s basic linker (Fig. 4A). We previously demonstrated that an overlapping region of the phosphorylated binding motif, Stu2^592-607^, is the Stu2 nuclear localization signal (Carrier *et al.*, 2022). Thus, we introduced mutations in a *STU2-NLS*-containing construct to evaluate the effects independently of any confounding effects on nuclear localization. Consistent with the hypothesis above, Stu2^S603A^-NLS-GFP, Stu2^Δ600-605^-NLS-GFP, and Stu2^Δ633-647^-NLS-GFP all showed more Stu2 at kinetochores than did Stu2^WT^-NLS-GFP in mitotically arrested cells (Fig. 4C). To test the second prediction, we inhibited Cdc28 activity using *cdc28-as1* and addition of 1NA-PP1 (Ubersax *et al.*, 2003). This experiment also showed higher levels of Stu2-NLS-GFP at kinetochores in *cdc28-as1* cells than in cells expressing *CDC28^WT^* (Fig. 4D), consistent with Cdc28-dependent priming of Stu2:Cdc5 association.

We also tested whether mutations in the conserved, putative polo-box binding motifs in Stu2 would disrupt its displacement from kinetochores by tethered Cdc5, using cells expressing Stu2^WT^-NLS-GFP, Stu2^Δ600-605^-NLS-GFP, and Stu2 ^Δ633-647^-NLS-GFP. Consistent with results in experiments described above, Stu2^WT^-NLS-GFP levels decreased at kinetochores when Cdc5-FRB was tethered to Nuf2-FKBP (Fig. 4E). Mutation of the basic patch, either to *stu2 ^Δ600-605^* or *stu2^Δ633-647^*, rendered Stu2 “resistant” to kinetochore-associated Cdc5 activity. These data strongly suggest that the basic patch residues Stu2^600-607^ and Stu2^633-647^ mediate the interaction between Stu2 and the Cdc5 polo-box domains in cells, and furthermore, that Stu2^T886^ is a direct target of Cdc5.

### Stu2 nuclear localization is regulated by Cdc28 phosphorylation of the conserved basic linker patch

The finding that Stu2^600-607^ is a polo-box binding motif for Cdc5 was somewhat surprising, as we had previously characterized this same region of Stu2 as its nuclear localization signal (Carrier *et al.*, 2022). Furthermore, our data suggest that S603 within this region is phosphorylated, likely by Cdc28, to facilitate Cdc5 binding. This observation prompted us to investigate whether S603 phosphorylation could also regulate Stu2 nuclear import in addition to priming an interaction with Cdc5 (Usui *et al.*, 2003; Ma *et al.*, 2007; Carrier *et al.*, 2022). As before, to show that this conserved basic patch region of Stu2 is sufficient to ensure nuclear localization, we fused these residues to a GFP-GST construct and monitored nuclear localization. In the absence of Stu2^592-607^, GFP-GST is diffuse throughout the cell; a construct containing Stu2^592-607^-GFP-GST accumulates in the nucleus (marked by the nuclear pore protein Nup2; Fig. 5A-B, see also (Carrier *et al.*, 2022)). Furthermore, the nuclear localization activity of Stu2’s basic linker is cell cycle-dependent. Cells arrested in G1 contain less nuclear Stu2^592-607^-GFP-GST than cells arrested in mitosis (Fig. 5B). The control SV40^NLS^-containing fusion does not exhibit cycle-dependent nuclear import, showing similarly high levels in both G1 and mitosis. Mutating residues required for importin α interaction (*stu2^K598A R599A^* i.e. “*KR/AA*”) (Carrier *et al.*, 2022) also disrupts this regulation, resulting in consistently low nuclear import across both cell cycle stages (Fig. 5B). Furthermore, an S603A mutation resulted in impaired NLS function of Stu2^592-607^-GFP-GST, while a phosphomimetic S603E partially rescued nuclear localization (Fig. 5C). We propose that the rescue was only partial because a charged amino acid in this case is a poor mimic of phosphorylated serine. We also observed lower mitotic nuclear accumulation of full length Stu2-GFP in *stu2^S603A^* cells than in *STU2^WT^* cells, showing that the same regulation also occurs in the full protein context (Fig. 5D).

**Figure 5.**
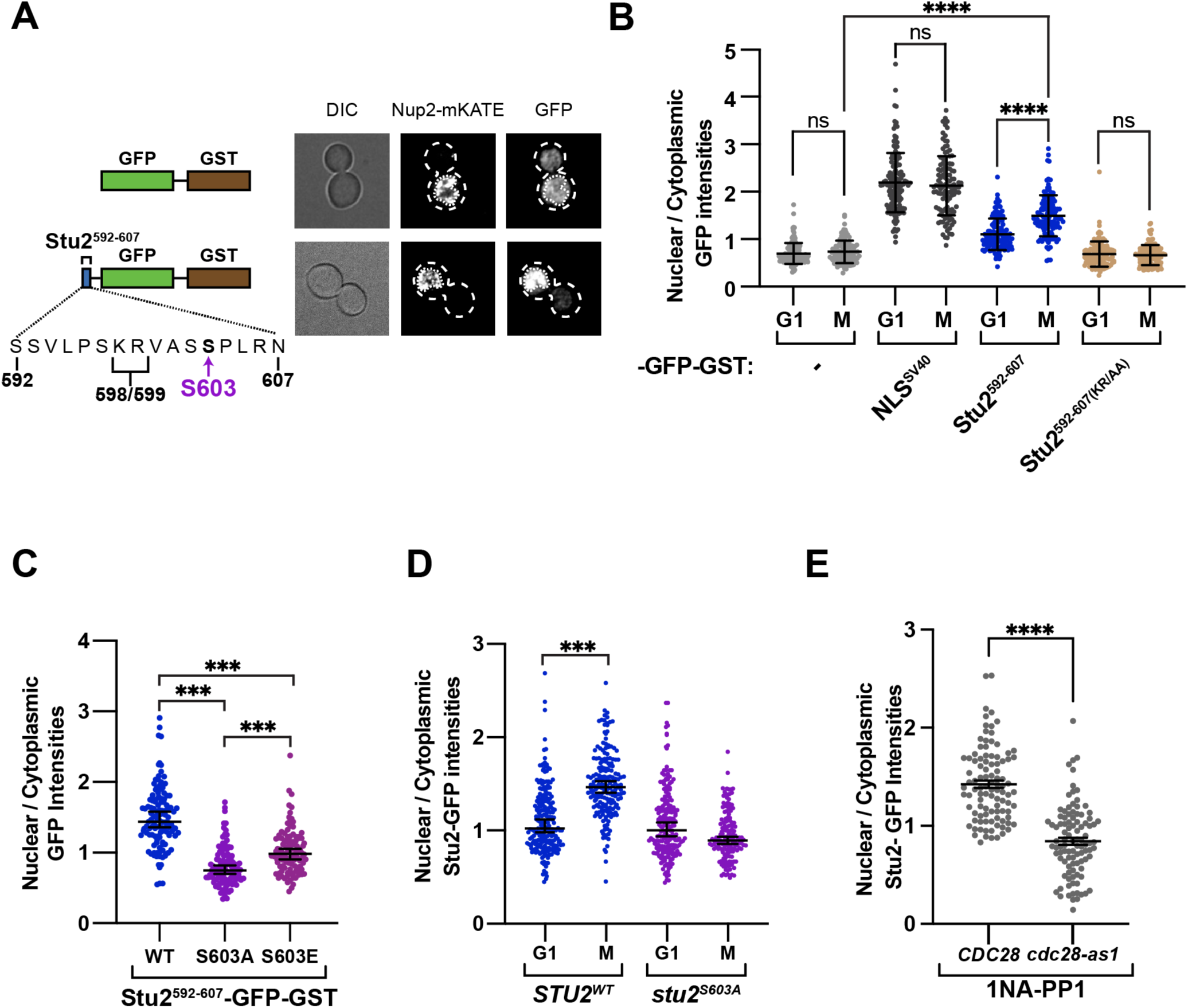
Phosphorylated Stu2^S603^ and Cdc28 activity are required for Stu2 nuclear import. A. Exponentially growing *stu2-AID cdc20-AID NUP2-mKate* cells ectopically expressing GFP-GST fused constructs (GFP-GST, M2390; Stu2^592-607^-GFP-GST, M2392) were treated with 500 µM auxin for 2 hours to arrest cells in metaphase. Cells were fixed and the Nup2-mKate and GFP signals were imaged. LEFT: Schematic of the constructs used. RIGHT: Representative images of DIC, Nup2-mKate, and GFP signals for each strain with an outlined nucleus. B. Exponentially growing *stu2-AID* cells ectopically expressing GFP-GST fused constructs (GFP-GST, M2441; NLS^SV40^-GFP-GST, M2442; Stu2^592-607^-GFP-GST, M2443; or Stu2^592-607(KR/AA)^-GFP-GST, M2439) as well as *NUP2-mKate* were treated with α-factor for 2.5 hours to arrest cells in G1 and then auxin for 30 minutes to degrade Stu2-AID. And exponentially growing *stu2-AID cdc20-AID* cells ectopically expressing GFP-GST fused constructs (GFP-GST, M2390; NLS^SV40^-GFP-GST, M2391; Stu2^592-607^-GFP-GST, M2392; or Stu2^592-607(KR/AA)^-GFP-GST, M2437) as well as *NUP2-mKate* were treated with auxin for 2 hours to degrade Stu2-AID and Cdc20-AID to arrest cells in metaphase. Cells were fixed and Nup2-mKATE and -GFP-GST construct signals were imaged. The ratios of nuclear to cytoplasmic GFP intensities were quantified. Each data point represents this ratio for a single cell. Mean and standard deviation are shown. n=107-139 cells; p values were determined using a two-tailed unpaired t test (**** = p<0.0001). C. Exponentially growing *stu2-AID cdc20-AID* cells ectopically expressing a GFP-GST fusion construct (Stu2^592-607^, M2392; Stu2^592-607(S603A)^, M2438; Stu2^592-607(S603E)^, M2726) were treated with auxin for 2 hours to degrade Stu2-AID and Cdc20-AID to arrest cells in metaphase. Cells were fixed and Nup2-mKATE and GFP signals were imaged. The ratios of nuclear to cytoplasmic GFP intensities were quantified. Each data point represents this ratio for a single cell. Bars represent median. Error bars are 95% confidence interval. n=104-124 cells; p-values were determined using a two-tailed unpaired test (*WT* vs *S603A*, p<0.0001; *WT* vs *S603E*, p<0.0001; *S603A* vs *S603E*, p<0.0001). D. Exponentially growing *stu2-AID* cells ectopically expressing full-length *STU2-GFP* alleles (*STU2^WT^-GFP*, M2298; or *stu2^S603A^-GFP*, M2351) were treated with alpha factor for 2.5 hours to arrest cells in G1 then with auxin for 30 minutes to degrade Stu2-AID, or cells as above but with *cdc20-AID* (*STU2^WT^-GFP,* M2208; or *stu2^S603A^-GFP,* M2217) were treated with auxin for 2 hours to degrade Stu2-AID and Cdc20-AID to arrest cells in metaphase. Cells were fixed and Nup2-mKATE and Stu2-GFP signals were imaged. The ratios of nuclear to cytoplasmic GFP intensities were quantified. Each data point represents this ratio for a single cell. Median and the 95% confidence interval are shown. n=100-165 cells; p-values were determined using a two-tailed unpaired t-test (*STU2^WT^* G1 vs. M, p<0.0001; *stu2^S603A^* G1 vs. M, p<0.0001). E. Exponentially growing *stu2-AID Nup2-mKate* cells with ectopic *STU2-GFP* and *CDC28^WT^* (M5762) or *cdc28-as1* (M5770) were treated with nocodazole for 2.5 hours to arrest in mitosis. Auxin and 1NA-PP1 were added 30 minutes prior to harvesting; cells were fixed and Nup2-mKate and GFP signals were imaged. The ratios of nuclear to cytoplasmic GFP intensities were quantified. Each data point represents this ratio for a single cell. Bars represent mean of n=100 individual measurements. Error bars are S.E.M. p-values were determined using a two-tailed unpaired t-test (*CDC28^WT^* vs *cdc28-as1*, p<0.0001).

Because our mutational analysis suggested that Cdc28 phosphorylation is required to drive Stu2 nuclear localization, we tested the importance of Cdc28 activity directly. We arrested *CDC28^WT^* and *cdc28-as1* cells in mitosis and added the inhibitor 1NA-PP1 (Ubersax *et al.*, 2003). In *cdc28-as1* cells, Stu2 nuclear accumulation in mitosis was lower than in *CDC28^WT^* cells, indicating that Cdc28 activity is required for proper Stu2 nuclear localization during mitosis (Fig. 5E). These results are consistent with Cdc28 regulation of Stu2 nuclear localization through phosphorylation of S603. They also indicate that the same region, and the same phosphorylated residue, facilitate both nuclear import and priming of Stu2:Cdc5 association.

### PP2A^Cdc55^ counteracts Cdc5 phosphorylation of Stu2 during metaphase

To examine the precise timing of Stu2 removal from kinetochores, we determined the Stu2 signal relative to Ndc80 in cells harboring *NDC80-mKate* and either *STU2^WT^-GFP* or *stu2^T866V^-GFP* after release from a G1 arrest, as previously described (Fig. 1A). To correlate any observed changes with cell cycle stage, we considered the ratio of Stu2:Ndc80 and the distance between the associated bilobed Ndc80 foci. In mitosis, Ndc80c distance is a close proxy of spindle length, with ‘short’ distances (< 1.5 µm) indicating pre-anaphase cells and ‘long’ distances (> 1.5 µm), anaphase cells (Joglekar, Bloom and Salmon, 2009; Marco *et al.*, 2013). We found no difference in Stu2:Ndc80c levels between *STU2^WT^-GFP* and *stu2^T866V^-GFP* cells before anaphase onset, either when Ndc80c appeared as a single, non-bilobed focus or with short (pre-anaphase) Ndc80c distance (Fig. 6A, Fig. S5A). In cells with longer (anaphase-like) Ndc80c distance, *STU2^WT^-GFP* cells had sharply lower Stu2:Ndc80c association than *stu2^T866V^-GFP* cells (Fig. 6A). This difference was consistent across cells at every “anaphase-like” Ndc80c distance, suggesting that phosphorylation of Stu2 T866 occurs rapidly at anaphase onset (Fig 6B, Fig. S5B).

**Figure 6.**
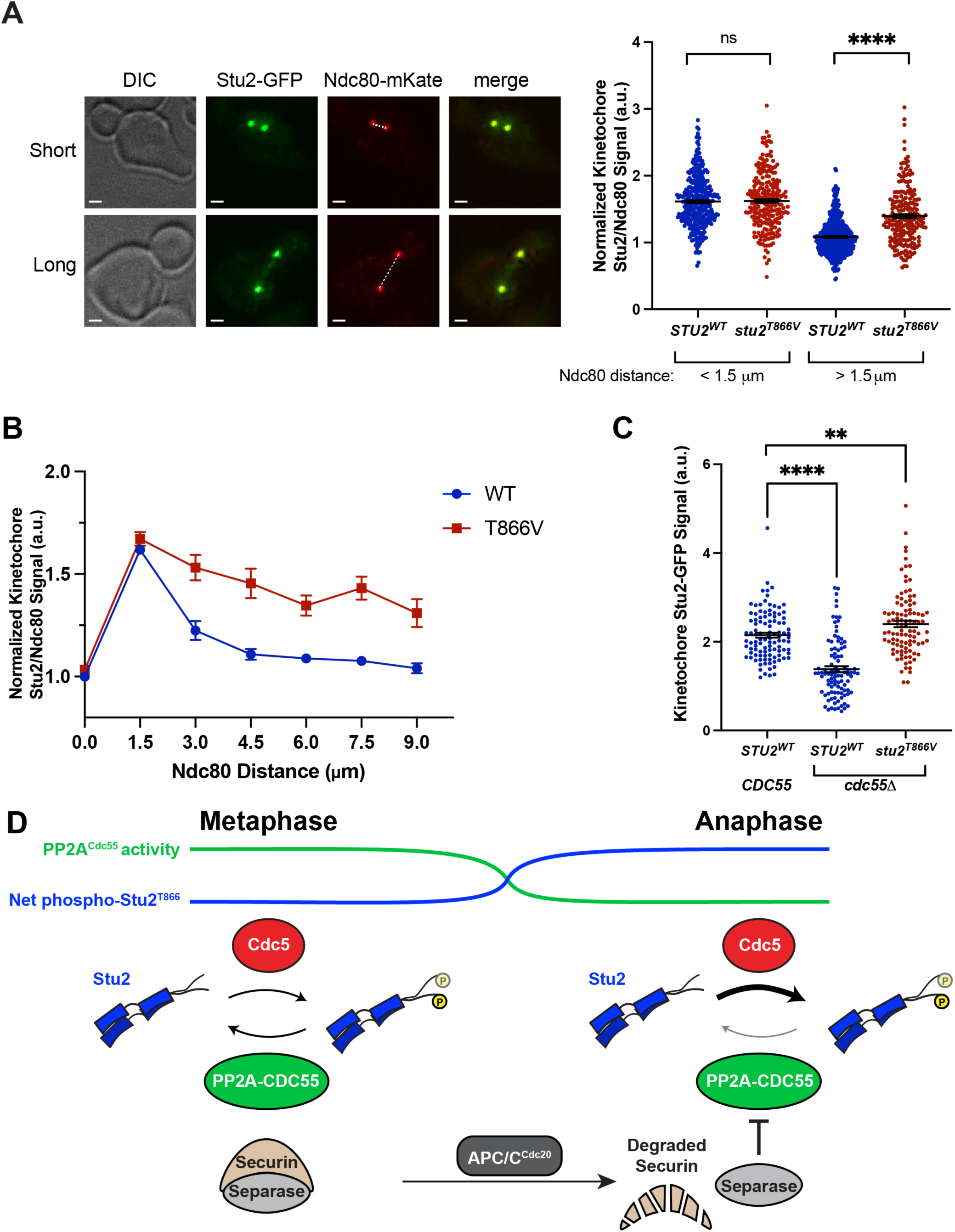
Stu2^T866^ phosphorylation is opposed by PP2A^Cdc55^ until anaphase onset. A. Exponentially growing *stu2-AID* cells expressing *STU2-GFP and NDC80-mKate* (M3774) or *stu2^T866V^-GFP* and *NDC80-mKate* (M4429), were released from a G1 arrest into auxin-containing media. Samples were taken every 15 minutes, fixed, and imaged. LEFT: Representative images of mitotic cells with pre-anaphase-like Ndc80 distance (Short, < 1.5 µm) and anaphase-like Ndc80 distance (Long, > 1.5 µm). Scale bars are 1 µm. Dotted line indicates how Ndc80 distance was measured. RIGHT: Kinetochore Stu2-GFP signal and Ndc80-mKate signal were measured for individual puncta. Ratio of Stu2/Ndc80c signal plotted for cells with Short (< 1.5 µm) or Long (> 1.5 µm) Ndc80 distance. Measurements are an average from 2 replicate time course experiments. Bars represent the mean of n=212-492 individual measurements. Error bars are S.E.M. p-value from unpaired t-tests (< 1.5 µm *STU2^WT^* vs *stu2^T866V^*, p=0.8375; > 1.5 µm *STU2^WT^* vs *stu2^T866V^*, p<0.0001). B. Exponentially growing *stu2-AID* cells expressing *STU2-GFP and NDC80-mKate* (M3774) or *stu2^T866V^-GFP* and *NDC80-mKate* (M4429), were cultured and imaged as in A. Ratio of Stu2/Ndc80c signal plotted for cells binned together by Ndc80 distance (0 µm: 0 µm ≤ x ≤1.5 µm; 1.5 µm: 1.5 µm < x ≤ 3 µm, etc.). Data points are mean of many individual measurements (*STU2^WT^*: 0 µm n=337, 1.5 µm n=345, 3.0 µm n=42, 4.5 µm n=111, 6.0 µm n=108, 7.5 µm n=104, 9.0 µm n=110. *stu2^T866V^*: 0 µm n=274, 1.5 µm n=182, 3.0 µm n=63, 4.5 µm n=48, 6.0 µm n=58, 7.5 µm n=50, 9.0 µm n=38). Error bars are S.E.M. C. Exponentially growing *stu2-AID cdc20-AID* cells with an ectopically expressed *STU2-GFP* allele (M5335) and *stu2-AID cdc20-AID cdc55Δ* cells with ectopic *STU2^WT^*-GFP (M5337) or *stu2^T866V^-GFP* (M5611) were cultured in auxin-containing media to arrest cells in metaphase. Cells were fixed and imaged to determine Kinetochore-Proximal Stu2-GFP. Bars represent the average of n=98-104 individual measurements. Error bars are S.E.M. p-values from two-tailed unpaired t-tests (*CDC55 STU2^WT^* vs *cdc55Δ STU2^WT^*, p<0.0001; *CDC55 STU2^WT^* vs *cdc55Δ stu2^T866V^*, p=0.0056). D. Model of PP2A^Cdc55^ opposing Cdc5 activity against Stu2^T866^. In metaphase (LEFT) PP2A^Cdc55^ is highly active and removes phosphorylation from Stu2^T866^, this cycling results in a steady state low level of pT866. In anaphase (RIGHT) separase inhibits PP2A^Cdc55^, allowing Cdc5 to “win” and resulting in rapid net phosphorylation of Stu2^T866^.

How do cells exert precise temporal control to achieve rapid phosphorylation of Stu2^T866^ at anaphase onset? Since specific upregulation of Cdc5 activity seemed unlikely, we considered the possibility that a counteracting mechanism to Cdc5 phosphorylation is abruptly inactivated. The activity of a phosphatase that counteracts Cdc5 phosphorylation could provide the “switchlike” response observed if such a phosphatase were inhibited precisely at anaphase onset. One major class of phosphatases active in mitosis are members of the protein phosphatase 2A (PP2A) family (Queralt *et al.*, 2006; Yellman and Burke, 2006; Clift, Bizzari and Marston, 2009; Yaakov, Thorn and Morgan, 2012; Moyano-Rodriguez and Queralt, 2019). These heterotrimeric phosphatases have distinct regulatory subunits, such as Cdc55 and Rts1, to specify the substrate to be dephosphorylated. If PP2A counteracted Cdc5 phosphorylation of Stu2^T866^ in metaphase, we would predict that depletion of the proper PP2A regulatory subunit would result in higher levels of pT866 and lower Stu2 signal at kinetochores. We therefore monitored the Stu2 kinetochore signal in metaphase-arrested cells with both *cdc55-AID* and *rts1-AID*. While the effect was smaller than that produced by the phosphomimetic *stu2^T866E^* allele, possibly due to inefficiency of these AID alleles (personal communication with Ethel Queralt), depletion of Cdc55-AID but not Rts1-AID led to a decrease in Stu2 kinetochore association (Fig. S5C). As a more robust test for whether PP2A^Cdc55^ affects Stu2^T866^ phosphorylation, we constructed yeast strains harboring a deletion of *CDC55* and measured Stu2-GFP levels in metaphase. Consistent with the idea that PP2A^Cdc55^ opposes Stu2^T866^ phosphorylation, *cdc55Δ* cells showed lower Stu2 levels at kinetochores than did wild-type cells; the effect depended on Stu2^T866^ phosphorylation, as *cdc55Δ stu2^T866V^* cells showed higher kinetochore-Stu2 levels than did wild-type cells (Fig. 6C). These results suggest that PP2A^Cdc55^ counters Cdc5 phosphorylation of Stu2^T866^ during metaphase.

If this model is correct, how is PP2A^Cdc55^ rapidly downregulated at anaphase onset? Prior genetic and phosphoproteomic studies suggest a plausible mechanism. They indicate that phosphatase PP2A^Cdc55^ counteracts general Cdc5 phosphorylation in metaphase (Touati *et al.*, 2019) and that threonine residues phosphorylated by Cdc5 appear to be more actively targeted. This counter-regulation by PP2A^Cdc55^ ends abruptly at the onset of anaphase, driven by the activation of separase, which inhibits PP2A^Cdc55^ in a proteolysis-independent manner (Queralt *et al.*, 2006; Calabria *et al.*, 2012; Touati *et al.*, 2019). Loss of PP2A activity allows Cdc5 to “win” and results in net phosphorylation of many Cdc5 substrates, including the mitotic exit regulator Net1 and likely also Stu2 (Fig. 6D). Our results are in good agreement with these prior studies and support the model of PP2A^Cdc55^ inhibition of Cdc5 activity through our extensive mutational analysis of a substrate, Stu2.

### Stu2^T866^ modification and PP2A^Cdc55^ activity are important for proper mitotic spindle maintenance

Finally, we sought to determine the cellular role of Stu2^T866^ phosphorylation. Stu2 has been previously implicated in anaphase spindle elongation (Severin *et al.*, 2001; Al-Bassam *et al.*, 2006). Because Stu2^T866^ phosphorylation appears to occur at the metaphase-anaphase transition, its relocation from the kinetochore may be crucial for anaphase spindle elongation. Disruption of this process could therefore lead to cellular defects. As previously observed (Usui *et al.*, 2003; Ma *et al.*, 2007), we saw Stu2-GFP signal at the spindle midzone (i.e. between the kinetochore-associated foci), which increases significantly at the onset of anaphase. These observations suggest that Stu2 relocates along interpolar microtubules once it is released from kinetochores (Fig. 6A, left panels). We investigated whether the increased kinetochore association of Stu2^T866V^ corresponded with reduced association to the spindle midzone and indeed found, in cells with elongated spindles, less midzone localized Stu2-GFP in *stu2^T866V^* cells than in *STU2^WT^* cells (Fig. 7A-B). There was no difference in the overall length of anaphase spindles between *STU2^WT^* and *stu2^T866V^* cells, showing that decreased Stu2 signal along microtubules was not due to differences in anaphase spindle length (Fig. 7C).

**Figure 7.**
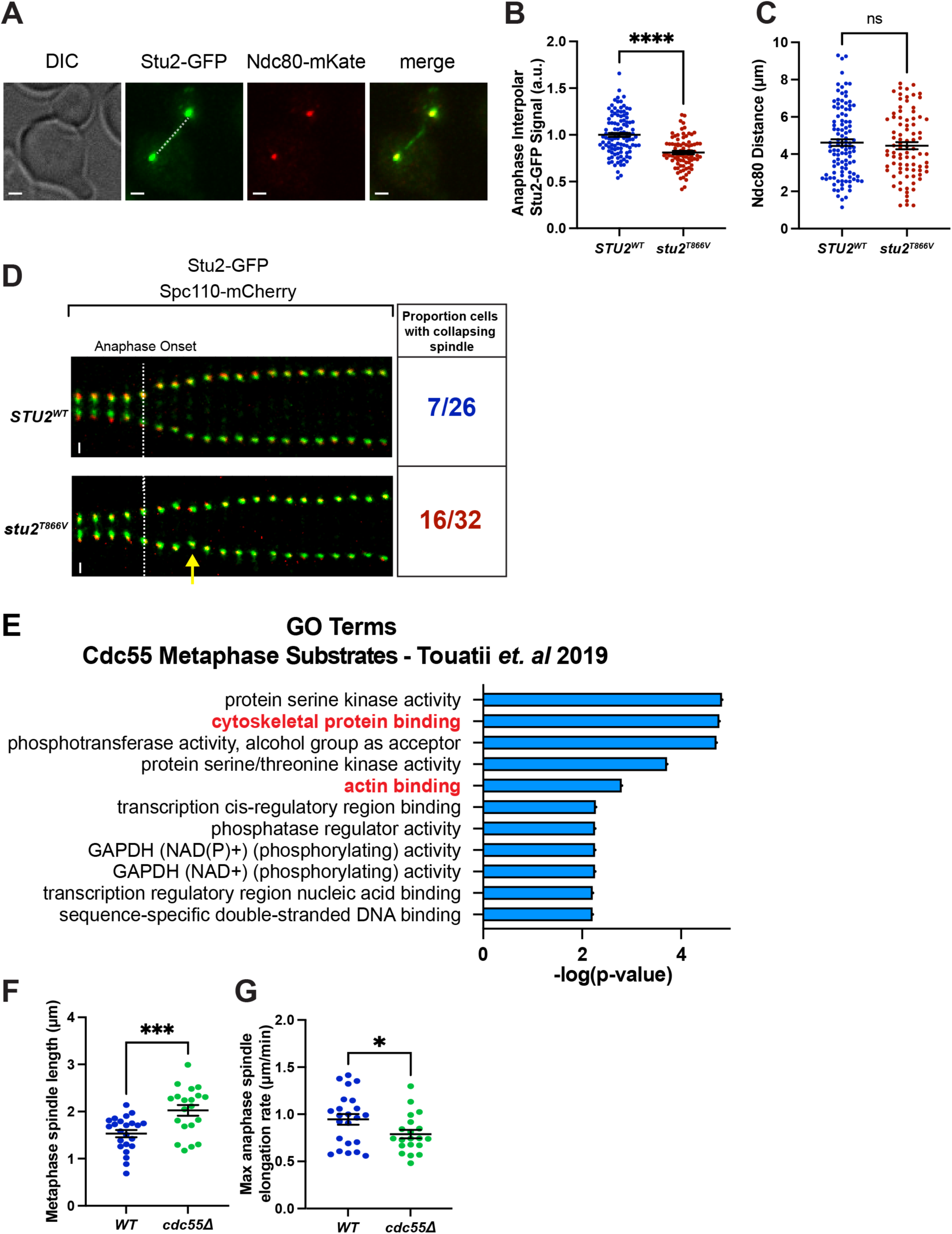
Stu2^T866^ phosphorylation is important for maintenance of anaphase spindle progression. A. Exponentially growing *stu2-AID* cells expressing *STU2-GFP and NDC80-mKate* (M3774) or *stu2^T866V^-GFP* and *NDC80-mKate* (M4429), were released from a G1 arrest into auxin-containing media. Samples were taken every 15 minutes, fixed, and imaged. Representative images of anaphase cells showing interpolar Stu2-GFP signal as dashed white line. Scale bars are 1 µm. B. Cells grown as in A. were imaged to determine interpolar Stu2-GFP signal for *STU2-GFP^WT^* for *stu2^T866V^-GFP* cells in anaphase. Bar represents mean of n=85-108 individual measurements. Error bar is S.E.M. p-value from two-tailed unpaired t-test (*STU2^WT^* vs *stu2^T866V^*, p<0.0001). C. Cells grown as in A. were imaged to determine distance between Ndc80 puncta for *STU2^WT^-GFP* and *stu2^T866V^-GFP* cells in anaphase. Bar represents mean of n=85-108 individual measurements. Error bar is S.E.M. p-value from two-tailed unpaired t-test (*STU2^WT^* vs *stu2^T866V^*, p=0.5418). D. Exponentially growing *stu2-AID, SPC110-mCherry* cells ectopically expressing *STU2-GFP* variants (*STU2^WT^-GFP*, M2429; *stu2^T866V^-GFP*, M5309) were treated with auxin for 30 minutes. Spc110-mCherry and Stu2-GFP signals were imaged every minute. Representative time course of a cell progressing through anaphase beginning 5 minutes prior to anaphase onset until 15 minutes post-anaphase onset. The white dotted line indicates anaphase onset. Arrow indicates instance of spindle collapse/regression. Proportion of cells that showed spindle regression phenotype indicated below time courses. E. Gene ontology analysis of 319 phophosites that were shown to be affected by Cdc55 activity in mitosis. See (Touati *et al.*, 2019). F. Exponentially growing *SPC110-mCherry MTW1-3GFP* cells containing *CDC55^WT^* (M1174) or *cdc55Δ* (M5848) we imaged every minute. Spindle length of cells in metaphase was determined by measuring the distance between Spc110-mCherry foci. Each data point represents an individual cell. Bars represent the average of n=20-23 individual measurements. Error bars are S.E.M. p-values from two tailed unpaired t-test (*CDC55^WT^* vs *cdc55Δ* p=0.0006). G. Cells grown and imaged as in F were analyzed to determine maximum rates of spindle elongation over a 2-minute period for each individual cell. Each data point represents a single cell. Bars represent the average of n=20-23 individual measurements. Error bars are S.E.M. p-values from two-tailed unpaired t-test (*CDC55^WT^* vs *cdc55Δ* p=0.0416).

To assess directly the effects of Stu2 localization on anaphase spindle elongation, we used live-cell imaging of *STU2^WT^-GFP* and *stu2^T866V^-GFP* cells, which also had spindle poles marked with *SPC110-mCherry.* We imaged cells every minute during mitosis and tracked the distance between Spc110 foci to measure mitotic spindle elongation. Both *STU2^WT^* and *stu2^T866V^* cells showed some frequency of transiently collapsing mitotic spindles under our imaging conditions, but spindle regression occurred almost twice as frequently in *stu2^T866V^* cells (7/26 *STU2*^WT^, 16/32 *stu2^T866V^*; Fig. 7D). Moreover, the maximum spindle elongation rate was lower in *stu2^T866V^* than in *STU2^WT^* cells (Fig. S5D). These observations suggest that Stu2^T866^ is phosphorylated at anaphase onset, causing a portion of the cellular Stu2 to relocalize to interpolar microtubules and regulate anaphase spindle elongation. This switchlike modification of Stu2 may be crucial for driving the rapid cytoskeletal reorganization required during anaphase.

Our observation that Stu2 appears to be regulated by the Cdc5-PP2A^Cdc55^ network led us to ask whether cells use this pathway to regulate other MAPs, to drive large microtubule cytoskeletal remodeling during anaphase. We performed a GO term analysis of 316 sites previously identified as regulated by PP2A^Cdc55^ during metaphase (Touati *et al.*, 2019). This analysis showed that many microtubule and actin cytoskeleton remodelers, including the MAPs Bim1 and Stu1, are among the metaphase targets of Cdc55 (Fig. 7E). This result is consistent with the idea that PP2A^Cdc55^ activity is important for regulating the cytoskeleton changes that happen at anaphase onset.

Our GO-term analysis of PP2A^Cdc55^ substrates, combined with mutational analysis of Stu2, led us to hypothesize that PP2A^Cdc55^ broadly regulates the mitotic spindle. To test this notion directly, we performed live-cell imaging of *CDC55^WT^* and *cdc55Δ* cells expressing *SPC110-mCherry* and *MTW1-GFP*, tracking the distance between Spc110 foci during mitosis. Consistent with our hypothesis, *cdc55Δ* cells had longer spindles in metaphase than *CDC55^WT^* and had lower maximum spindle elongation rates in anaphase (Fig. 7F-G). From our Stu2 mutational analysis, we expected a spindle defect in metaphase, because PP2A^Cdc55^ actively counteracts Cdc5 phosphorylation at that stage. The persistence of spindle defects into anaphase suggests that the precise timing of phosphorylation events regulated by Cdc5-PP2A^Cdc55^ is critical for spindle function. Overall, these findings are in line with the notion that PP2A^Cdc55^ activity supports mitotic spindle maintenance, likely through post-translational modification of Stu2 and other microtubule-associated substrates (Fig. S5E-G).

## DISCUSSION

The XMAP215 family member Stu2 has multiple cellular roles and functions at microtubule organizing centers, kinetochores, and microtubules during different points in the cell cycle (van Breugel, Drechsel and Hyman, 2003; Al-Bassam *et al.*, 2006; Hsu and Toda, 2011; Miller, Asbury and Biggins, 2016; Zahm *et al.*, 2021). We have studied their regulation, including the import into the nucleus and modulation of kinetochore and microtubule localization. In particular, we have investigated how phosphorylation of Stu2 at two specific sites governs its subcellular activities. We find that Cdk/Cdc28 activity facilitates nuclear import of Stu2, through phosphorylation of Stu2^S603^ in the nuclear localization sequence in Stu2’s basic linker. We further find that phosphorylation of a threonine in Stu2’s CTS (Stu2^T866^) reduces association of Stu2 with Ndc80c. Stu2^T866^ phosphorylation depends on the polo-like kinase, Cdc5, and Cdc28 phosphorylation of S603 primes the Stu2:Cdc5 interaction. PP2A^Cdc55^ opposes Stu2^T866^ phosphorylation in metaphase. Upon inhibition by the newly activated separase, downregulation of PP2A^Cdc55^ leads to rapid accumulation of phosphorylated Stu2^T866^ at anaphase onset (Fig. 8) (Queralt *et al.*, 2006). Modification of this site allows cells to relocalize a pool of Stu2 to the spindle midzone during anaphase for maintenance of anaphase spindle stability. This regulatory event is likely one of many modifications that the Cdc5-PP2A^Cdc55^ network catalyzes to elicit the rapid cytoskeletal changes required for entry into anaphase (Touati *et al.*, 2019).

**Figure 8.**
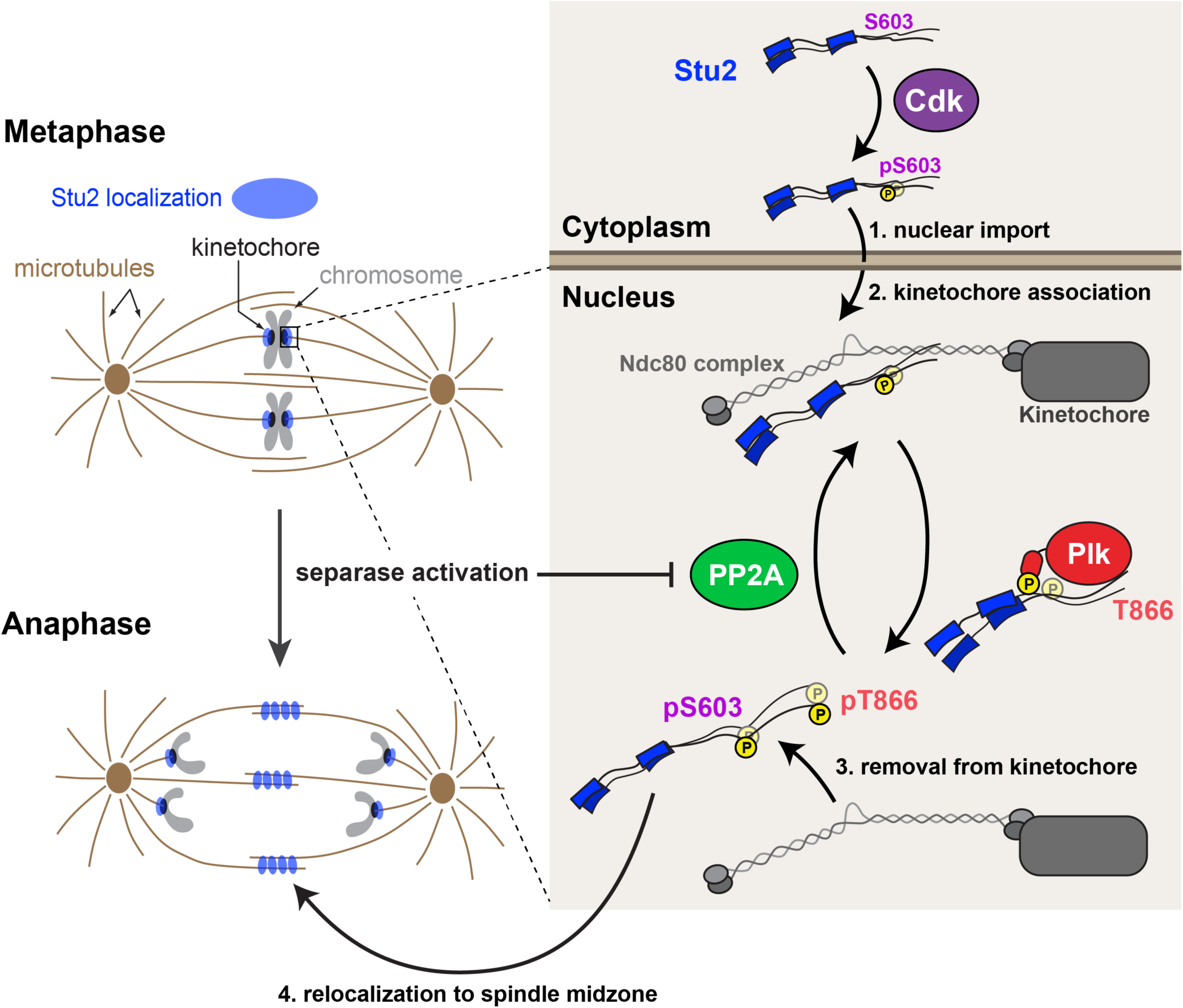
Model of dynamic kinase and phosphatase regulation of Stu2 kinetochore localization and function. In G1 phase, Stu2 is predominantly localized to the cytoplasm. As Cdk activity increases in the cell cycle, Stu2^S603^ becomes phosphorylated, and Stu2 is imported into the nucleus. In the nucleus, Stu2 binds to Ndc80c, and carries out a required function for chromosome biorientation. During metaphase, Plk1\Cdc5 interacts with Stu2 via binding phosphorylated Stu2^S603^. Plk1 phosphorylates Stu2^T866^ and PP2A^Cdc55^ opposes this phosphorylation. At anaphase onset, PP2A^Cdc55^ activity is inhibited by separase and Stu2^T866^ becomes predominantly phosphorylated. This leads to reduced Stu2:Ndc80c association, and relocalization of some Stu2 to spindle microtubules to regulate anaphase spindle elongation.

The rapid phosphorylation of Stu2^T866^ at anaphase onset could serve other cellular purposes. One mechanism we considered is whether Stu2-dependent kinetochore functions are affected by Stu2^T866^ phosphorylation, including Stu2’s proposed role in error correction (Miller, Asbury and Biggins, 2016; Miller *et al.*, 2019). Kinetochore error-correction mechanisms are thought to be downregulated at anaphase onset to address the “anaphase problem” (Vázquez-Novelle *et al.*, 2010) as kinetochore-microtubule attachments are under low tension during anaphase. Ipl1, a main effector of error-correction, is relocalized from centromeres to the midzone at anaphase onset through Cdc14-mediated dephosphorylation of Sli15 (Buvelot *et al.*, 2003; Khmelinskii *et al.*, 2007; Zimniak *et al.*, 2012; Cairo *et al.*, 2023). It is attractive to propose that phosphorylation of Stu2^T866^ could serve as a mechanism to control Stu2’s putative error-correction activity, but our data suggest that Stu2^T866^ modification is not the sole regulator of Stu2-dependent kinetochore function. The phenotypes observed in *stu2^T866V^* cells are not consistent with hyper-activated error correction, such as tethering Ipl1 to outer kinetochore substrates (Li, Garcia-Rodriguez and Tanaka, 2023), which compromises cell viability. Instead, it is more likely that Stu2^T866^ phosphorylation plays a role in redistributing Stu2 between kinetochores and microtubules to maintain the anaphase spindle, as our data show. Other protein-protein interactions mediated by Stu2’s C-terminus may also be regulated by Stu2^T866^ phosphorylation (Figure S5E-G) (Stangier *et al.*, 2018), similar to the regulation we have shown for Stu2:Ndc80c. Such additional controls may be necessary for spindle maintenance or other as-yet-undetermined functions. Our results also do not exclude the possibility that other post-translational modifications of Stu2 or Ndc80c may influence their association and/or kinetochore function (Greenlee *et al.*, 2022).

Our work provides new, detailed information concerning mechanisms for specifying substrates of Plk1/Cdc5. Many Cdc5 substrates require priming through proline-directed phosphorylation before they can interact with Cdc5 polo-box domain (Elia *et al.*, 2003), while others depend on hydrophobic interactions between the substrate and a distal face of the Cdc5 polo-box domain, separate from the canonical phosphopeptide binding region (Almawi *et al.*, 2020). A recent report also suggests a different mode of human Plk1:substrate interaction, mediated by electrostatic interactions on the canonical phosphopeptide binding surface of Plk1, but independent of substrate phosphorylation (Conti *et al.*, 2024). This interaction has been shown to be important for Plk1 to assist in nucleosome deposition in G1, a time at which Cdk activity is very low (reviewed in (Bloom and Cross, 2007)). Our results show that Stu2 has two polo-box interacting motifs: one dependent on phosphorylation of a Cdk/Cdc28 consensus site (Stu2^S603^) and another driven by hydrophobic interactions, both of which facilitate the Stu2:Cdc5 association. An important direction for future research will be to determine whether both interaction motifs are always used or if they are specifically engaged during different stages of the cell cycle, when Cdk activity levels vary. We also note that a previous study has suggested that Stu2 is a Cdc5 substrate, findings that nicely complement our mechanistic work here (Park *et al.*, 2008).

Furthermore, our results offer in-depth mutational analysis of a kinase-phosphatase network that controls the timing of many cellular events at anaphase onset. We show a new Cdc5-PP2A^Cdc55^ substrate that behaves like the prototypical Cdc5-PP2A^Cdc55^ substrate Net1 (Queralt *et al.*, 2006; Touati *et al.*, 2019). Phosphoproteomics experiments also showed that Cdc5 and PP2A^Cdc55^ modify many microtubule- and actin-cytoskeleton associated factors during the metaphase-anaphase transition (Fig. 7E, see also (Touati *et al.*, 2019)). More targeted studies show that PP2A^Cdc55^ regulates the actin cytoskeleton during the cell cycle (Jonasson *et al.*, 2016; Moyano-Rodríguez *et al.*, 2022). Complementing these findings, we show that PP2A^Cdc55^ also regulates the microtubule cytoskeleton, in part by controlling Stu2 localization. Additional microtubule regulators such as Bim1 or Stu1, which are also PP2A^Cdc55^ targets (Touati *et al.*, 2019), likely contribute to the large changes in the cytoskeleton during anaphase. Further work is needed to investigate the effects of Cdc5 and PP2A^Cdc55^ on these additional cytoskeleton regulatory proteins. We believe that misregulation of these factors may account for the spindle defects observed in cells induced to enter anaphase via TEV-Scc1 cleavage in the absence of separase activity (Uhlmann *et al.*, 2000). Prior phosphoproteomics studies of PP2A^Cdc55^ substrates did not detect Stu2^pT866^, suggesting that other important substrates in this pathway may have also escaped detection (Baro *et al.*, 2018; Touati *et al.*, 2019). In particular, several kinetochore, motor, and MAP proteins are predicted to be Cdc5 substrates, including Ndc80, Spc24, Kar3, Bim1, Sli15, and Cse4 (Ólafsson and Thorpe, 2015, 2020). Whether the Cdc5-PP2A^Cdc55^ network also regulates these or other proteins at anaphase onset could be addressed by the kinase tethering tools and mutational analyses described here. These issues remain an important avenue for future work.

Finally, there are reports from studies on human cells indicating anaphase-specific phosphorylation of Plk1 targets, particularly the microtubule regulator Prc1. These studies show that Prc1^T602^ is phosphorylated by Plk1 in metaphase, and that phosphorylated T602 accumulates rapidly in anaphase to facilitate proper function of the spindle midzone (Neef *et al.*, 2007; Hu *et al.*, 2012; Holder, Mohammed and Barr, 2020; Lim *et al.*, 2024). This pattern is reminiscent of the behavior of Stu2^T866^ and other Cdc5-PP2A^Cdc55^ threonine substrates in yeast. Furthermore, evidence suggests a similar inactivation of human PP2A^B55^ dephosphorylation during anaphase (Játiva *et al.*, 2019). The mechanisms described here may therefore represent a conserved strategy to enhance phosphorylation of Plk1 substrates rapidly at anaphase onset.

## METHODS

### Strain construction and microbial techniques

#### Yeast strains

*Saccharomyces cerevisiae* strains used in this study, all derivatives of M3 (W303), are described in Table S1. Standard media and microbial techniques were used (Sherman *et al.*, 1974). Yeast strains were constructed by standard genetic techniques. Construction of *DSN1-6His-3Flag* is described in (Akiyoshi *et al.*, 2010), *STU2-3FLAG* and *stu2-3V5-IAA7* in (Miller, Asbury and Biggins, 2016) and *TOR1-1*, *fpr1Δ*, and *MPS1-FRB:KanMX* in (Haruki, Nishikawa and Laemmli, 2008; Aravamudhan, Goldfarb and Joglekar, 2015). Construction of *MTWI-3GFP* is described in (Pinsky *et al.*, 2006). *NUF2-FKBP12:HisMX* construction was described in (Zahm *et al.*, 2021) *CDC5-FRB:KanMX*, *BUB1-FRB:KanMX*, and *IPL1-FRB:KanMX*, *cdc55-3HA-IAA7*, *rts1-3HA-IAA7*, *NDC80-mKate*, *BIK1-3FLAG* were constructed by PCR-based methods (Longtine *et al.*, 1998). Strains containing the previously described *pMET-CDC20* allele were provided by Frank Uhlmann. *CDC20-AID* and *cdc55Δ* containing strains were gifts from Adèle Marston. The *cdc5-1* allele was provided by David Morgan. For Quantitative Chromosome Transmission Fidelity (qCTF), strains with *MFA1-3XGFP* and the *(CEN3.L.YA5.1)MATalpha:LEU2* mini chromosome were generously provided by Rong Li. *SPC110-mCherry* containing strains were provided by Trisha Davis.

#### Plasmid construction

*pGPD1-TIR1* integration plasmids (pM76 for integration at *HIS3* or pM78 for integration at *TRP1*) were provided by Leon Chan. Construction of a 3HA-IAA7 tagging plasmid (pM69) as well as a *LEU2* integrating plasmid containing wild-type *pSTU2-STU2-3V5* (pM225) and *pSTU2-stu2Δ855– 888-3V5* (pM267) are described in (Miller, Asbury and Biggins, 2016; Miller *et al.*, 2019). Plasmid construction for *STU2-NLS-GFP* (pM659), Stu2^592-607^-GFP-GST (pM774), *STU2-GFP-GST* (pM772) as well as mutants of these plasmids, Stu2^592-607(S603A)^-GFP-GST (pM1362) and Stu2^592-607(S603E)^-GFP-GST (pM1410), are described in (Carrier *et al.*, 2022). *stu2^L869E,I873E,M876E^-3V5* construction is described in (Zahm *et al.*, 2021). *STU2-GFP* (pM488) and *STU2-3HA* (pM227) plasmids for this study were constructed by megaprimer mutagenesis as described in (Liu and Naismith, 2008; Tseng *et al.*, 2008). *STU2* variants were constructed by mutagenizing the above plasmids. Primers used in the construction of the above plasmids are listed in Table S2, and further details of plasmid construction including plasmid maps are available upon request.

#### Yeast Culture

Standard culture conditions for *Saccharomyces cerevisiae* (W303) were followed for all experiments unless otherwise indicated. Strains were grown to be in logarithmic growth phase for all experiments unless otherwise noted. Strains containing the temperature sensitive allele *cdc5-1* were grown at the permissive temperature of 23°C to maintain cell viability, then shifted to 37°C for 60 minutes to inhibit *cdc5-1*. For Cdk inhibition, strains harboring the analogue sensitive *cdc28-as1* allele were treated with 2.5 µM 1NA-PP1 for 30 minutes. To deplete Cdc20-AID for metaphase arrests IAA7 treatment was used as described below for 2.5 hours. For metaphase arrests using *pMET-CDC20* cells were arrested in rich media containing 8 mM methionine for 2.5 hours. To arrest cells in mitosis using nocodazole cells were treated with 10 μg/mL nocodazole for 2.5 hours. Alpha factor arrests were performed by treating cells with 10 μg/mL alpha factor for 3.5 hours.

#### Auxin inducible degradation

The AID system was used essentially as described (Nishimura *et al.*, 2009). Briefly, cells expressed C-terminal fusions of the protein of interest to an auxin responsive protein (IAA7) at the endogenous locus. Cells also expressed *TIR1*, which is required for auxin-induced degradation. 500 μM IAA (indole-3-acetic acid dissolved in DMSO; Sigma) was added to media to induce degradation of the AID-tagged protein. Auxin was added for 30 min prior to harvesting cells or as indicated in figure legends.

#### Spotting assay

For the spotting assay, the desired strains were grown for 2 days on plates containing yeast extract peptone plus 2% glucose (YPAD) medium. Cells were then resuspended to OD600 ∼1.0 from which a serial 1:5 dilution series was made and spotted on YPAD+DMSO, YPAD+500 μM IAA (indole-3-acetic acid dissolved in DMSO) or plates containing 3.5–5.0 μg/mL benomyl or 0.05 μg/mL rapamycin as indicated. Plates were incubated at 23°C for 2–3 days unless otherwise noted.

#### Quantitative Chromosome Transmission Fidelity

Chromosome Transmission Fidelity assay was carried out as described in (Zhu *et al.*, 2015)). Briefly, exponentially growing *stu2-AID* strains containing *MFA1-3XGFP* and *MC(CEN3.L.YA5.1)MATalpha:LEU2* and *STU2-3HA* variants were diluted into synthetic complete media containing auxin and grown for 16 hours. Loss of the mini-chromosome fragment was observed by measuring GFP signal by flow cytometry and quantified for 10,000 cells.

#### FRB/FKBP tethering

For re-tethering in culture, exponentially growing cultures were treated with 500 μM auxin and 0.20 μg/mL (200 ng/mL rapamycin) 30 min prior to harvesting. For spotting assays on plates, 0.05 μg/mL rapamycin was used.

#### Fluorescence microscopy

For imaging of fixed cells, cells were treated with 3.5% Formaldehyde in Kpi (e.g. 0.1 M potassium phosphate) buffer for 5 minutes. Cell images were collected with a DeltaVison Elite wide-field microscope system (GE Healthcare) equipped with a scientific CMOS camera, using 60X objective (Olympus; NA = 1.42 PlanApoN) and immersion oil with a refractive index of n = 1.516. A Z-stack was acquired over a 3 µm width with 0.2 µm Z-intervals. Images were deconvolved using the DeltaVision algorithm, maximally projected, and analyzed using the Fiji image processing package (ImageJ). Intensity of whole-cell Stu2-GFP, and Stu2-GFP and Ndc80-mKate kinetochore puncta was determined using Fiji. For live cell imaging, exponentially growing cultures, grown in synthetic complete media, were treated with 500 μM auxin 30 min prior to imaging to degrade Stu2-AID and analyzed for Stu2-GFP distribution and spindle length or left untreated to observe Mtw1-GFP and spindle length. For each strain, an aliquot of cells was pelleted and resuspended in a volume of synthetic complete media with 500 μM auxin to optimize cell density for imaging (OD600≈5). Cells were adhered to a coverslip coated in Concanavalin A as described in (Fees, Estrem and Moore, 2017) and the chamber was sealed using petroleum jelly. Cells were imaged using a DeltaVision Ultra as above. Images of the Spc110-mCherry signal, Stu2-GFP signal and DIC were acquired through the thickness of the cells using Z-stacks 0.3 µM apart. These images were acquired every 1 minute. All frames were deconvolved using standard settings. Image stacks were maximally projected for analysis of spindle lengths and Stu2 distribution. softWoRx image processing software was used for image acquisition and processing. Projected images were imported into FIJI for analysis. Distance between Spc110-mCherry puncta were measured every minute beginning 5 minutes prior to anaphase onset until 15 minutes post-anaphase onset. Anaphase onset was determined for a given cell by selecting the spindle length at the time point prior to which an increase in spindle length of at least 0.2 µM was observed and followed by an increase in spindle length over the next 3 time points. Spindle collapse events were defined as events of spindle regression measuring 0.2 µM or more between images. Maximum rate of spindle elongation for a given cell was calculated by determining the maximum difference in spindle length over a 2-minute time period and calculating the rate of spindle elongation over that time period. Metaphase spindle length was determine as the average spindle length of the five time points prior to anaphase onset.

#### Gene Ontology Term analysis

Genes for Gene Ontology analysis were compiled from mass spectrometry data of phosphosites that changed in the presence of *cdc55Δ* (See (Touati *et al.*, 2019), Figure 3G, also Supplemental Data S2). GO Terms determined as described in (Ashburner *et al.*, 2000) using The Gene Ontology Reference Server.

#### Multiple Sequence Alignment

Fungal proteins related to *Saccharomyces cerevisiae* Stu2 were identified using a PSI-BLAST (Altschul, 1997) search on NCBI. Multiple sequence alignments of the entire proteins were generated with ClustalOmega default parameters and displayed in JalView 1.8

#### Structure Prediction and Modeling

AlphaFold Server was used to produce AlphaFold 3 structural predictions of indicated protein complexes containing phosphorylated and un-phosphorylated polypeptides (Abramson *et al.*, 2024). Protein structures were modeled and compared in UCSF Chimera and ChimeraX. Building phosphates onto threonine in structure models was accomplished using ChimeraX.

### Protein biochemistry

#### Purification of native yeast protein

Native kinetochore particles were purified from asynchronously growing *S. cerevisiae* cells as described below. Dsn1-6His-3Flag was immunoprecipitated with anti-Flag essentially as described in (Akiyoshi *et al.*, 2010). Tagged yeast proteins (Bik1-3Flag, Stu2-3Flag) were similarly purified from yeast as follows. Cells were grown in yeast peptone dextrose (YPAD) rich medium. For strains containing *stu2-AID*, cells were treated with 500 μM auxin 30 min prior to harvesting. Protein lysates were prepared by mechanical disruption in the presence of lysis buffer using glass beads and a beadbeater (Biospec Products) or freezer mill (Spex Sample Prep). Lysed cells were resuspended in buffer H (BH; 25 mM HEPES pH 8.0, 2 mM MgCl_2_, 0.1 mM EDTA, 0.5 mM EGTA, 0.1% NP-40, 15% glycerol with 150 mM KCl) containing protease inhibitors at 20 μg/mL final concentration each for leupeptin, pepstatin A, chymostatin, and 200 μM phenylmethylsulfonyl fluoride and phosphatase inhibitors (0.1 mM Na-orthovanadate, 0.2 μM microcystin, 2 mM β-glycerophosphate, 1 mM Na pyrophosphate, 5 mM NaF) followed by centrifugation at 16,100 g for 30 min at 4°C to clarify lysate. Stu2-FLAG lysates were processed by centrifugation at 24,000 g for 90 min at 4°C to clarify the lysate. Dynabeads conjugated with anti-Flag or anti-V5 antibodies were incubated with extract for 3 hours with constant rotation, followed by three washes with BH containing protease inhibitors, phosphatase inhibitors, 2 mM dithiothreitol (DTT), and 150 mM KCl. Beads were further washed twice with BH containing 150 mM KCl and protease inhibitors. Associated proteins were eluted from the beads by boiling in 2X SDS sample buffer. Stu2-3Flag protein for mass spec analysis was eluted from beads with rapigest.

#### Immunoblot analysis

For immunoblot analysis, cell lysates were prepared as described above or by pulverizing cells with glass beads in sodium dodecyl sulfate (SDS) buffer using a bead-beater (Biospec Products). Standard procedures for sodium dodecyl sulfate-polyacrylamide gel electrophoresis (SDS-PAGE) and immunoblotting were followed as described in (Towbin, Staehelin and Gordon, 1979; Burnette, 1981). A nitrocellulose membrane (Bio-Rad) was used to transfer proteins from polyacrylamide gels. Commercial antibodies used for immunoblotting were as follows: α-Flag, M2 (Sigma-Aldrich) 1:3000; α-V5 (Invitrogen) 1:5000. Antibodies to Ndc80 (OD4) were a kind gift from Arshad Desai and used at 1:10,000. The secondary antibodies used were a sheep anti-mouse antibody conjugated to horseradish peroxidase (HRP) (GE Biosciences) at a 1:10,000 dilution or a donkey anti-rabbit antibody conjugated to HRP (GE Biosciences) at a 1:10,000 dilution. Antibodies were detected using the SuperSignal West Dura Chemiluminescent Substrate (Thermo Scientific).

#### Mass Spectrometry

Rapigest-eluted Stu2-3Flag protein was subjected to trypsin digestion prior to LC/MS -MS analysis. LC-MS/MS was performed on an Easy-nLC 1000 (Thermo Scientific) coupled to an LTQ-Orbitrap Elite mass spectrometer (Thermo Scientific) operated in positive ion mode. The LC system consisted of a fused-silica nanospray needle (PicoTip™ emitter, 50 µm ID x 20 cm, New Objective) packed in-house with Magic C18-AQ, 5mm and a trap (IntegraFrit™ Capillary, 100 µm ID x 2 cm, New Objective) containing the same resin as in the analytical column with mobile phases of 0.1% formic acid (FA) in water (buffer A) and 0.1% FA in acetonitrile (MeCN) (buffer B). The peptide sample was diluted in 20 µL of 0.1% FA, 2% MeCN and 8 µL was loaded onto the column and separated over 81 minutes at a flow rate of 300 nL/min with a gradient from 5 to 7% B for 2 min, 7 to 35% B for 60 min, 35 to 50% B for 1 min, hold 50% B for 8 min, 50 to 95% B for 1min, hold 95% B for 9 min. A spray voltage of 2500 V was applied to the nanospray tip. MS/MS analysis was performed for 80 minutes and consisted of 1 full scan MS from 400-1800 m/z at resolution 240,000 followed by data dependent MS/MS scans using 35% normalized collision energy of the 20 most abundant ions. Selected ions were dynamically excluded for 30 seconds. Raw MS/MS spectra from the analysis were searched against a W303 yeast strain protein database (downloaded 6/9/2022) containing common contaminants using Proteome Discoverer v3.1, with tryptic enzyme constraint set for up to two missed cleavages, oxidized methionine and phosphorylated serine, threonine and tyrosine set as a variable modification, carbamidomethylated cysteine set as a static modification and peptide MH+ mass tolerances set at 10 ppm. The peptide FDR was set at ≤1%.

#### Recombinant protein expression and purification

Ndc80dwarf protein was expressed and purified as described in (Valverde, Ingram and Harrison, 2016; Zahm *et al.*, 2021).

#### Fluorescence polarization binding assay

A Stu2 CTS peptide (855-888), a CTS peptide containing a phosphothreonine at position 866, and a CTS peptide with a randomized sequence, all containing a C-terminal cysteine residue and synthesized by the Tufts University Core Facility, were resuspended in a volume of buffer containing 100 mM Hepes pH 7.5, 100 mM NaCl, 0.5 mM TCEP (resuspension buffer) sufficient to yield a 2 mM peptide concentration. For labeling, the Stu2 CTS peptide was diluted to 100 µM in resuspension buffer to a final volume of 7 mL. To this solution was added 1 mL of 10 mM Oregon Green maleimide dissolved in DMSO. After a 24-hour incubation at 4°C, the labeled peptide was separated from unreacted dye by cation exchange chromatography with Source 15S resin (Cytiva). For the competition experiment, the peptides were subjected to serial 2-fold dilutions in resuspension buffer, and each dilution was mixed 1:1 with a solution containing 30 µM Ndc80dwarf, and 200 nM Oregon green-labeled CTS peptide, also in resuspension buffer. Fluorescence polarization was measured in triplicate in 96-well plates with an EnVision multi-mode plate reader (Perkin Elmer) located in the ICCB-Longwood Screening Facility at Harvard Medical School.

## Data Availability

The mass spectrometry data underlying Fig. 1B and Fig. S1B is available in supplemental material. Any other data are available from the corresponding author on reasonable request.

## Acknowledgments

We thank Arshad Desai for providing antibodies, Sue Biggins, David Morgan, Frank Uhlmann, Adèle Marston, Rong Li, Trisha Davis, and Leon Chan for strains and/or plasmids. We would like to thank Sue Biggins and members of the Miller lab for critical reading of the manuscript. We also thank the University of Utah Cell Imaging Core for maintaining the Delta Vision microscope facility. This work was supported in part by the Proteomics & Metabolomics Core Shared Resource of the Fred Hutch/University of Washington Cancer Consortium (P30 CA015704) as well as NIH Grants F31CA2717405 (to M.G.S.) and 5T32HG008962-05 (to M.G.S.), American Cancer Society Postdoctoral Fellowship PF-21-188-01-CCB (to J.A.Z), and 5 For the Fight (to M.P.M.), Pew Biomedical Scholars (to M.P.M.), and NIH grant R35GM142749 (to M.P.M.). S.C.H. is an investigator of the Howard Hughes Medical Institute.

## Author Contributions

M.G.S. & M.P.M. conceptualized the studies. M.G.S. performed the experiments and data analysis unless otherwise noted here. J.S.C. contributed data for Fig. 5A-D, and J.A.Z. contributed data for Fig. 2D. M.G.S. & M.P.M. wrote and edited the manuscript with assistance from S.C.H.

## SUPPLEMENTAL INFORMATION

**Figure S1.**
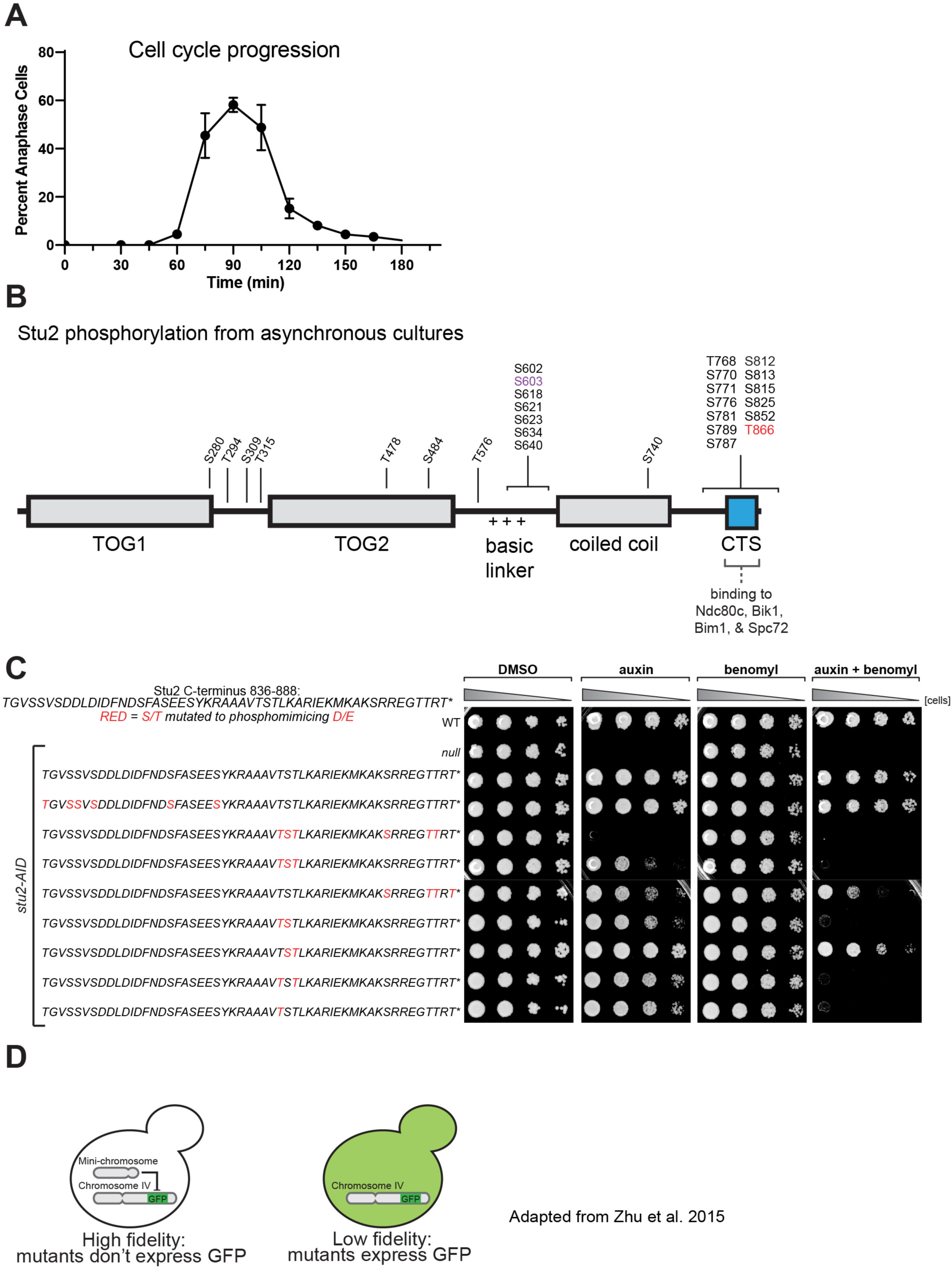
Stu2 phosphorylation and mutant viability phenotypes. A. Exponentially growing *stu2-AID* cells expressing *STU2-GFP and NDC80-mKate* (M3774), were released from a G1 arrest into auxin-containing media. Samples were taken every 15 minutes, fixed, and imaged as in Figure 1A. Percent large-budded cells were plotted for each time point. Data points are mean from 2 biological replicates and n=92-118 individual measurements. Error bars are S.E.M. B. Exponentially growing *STU2-3FLAG* (M498) cultures were harvested, lysed to produce protein sample, subjected to α-Flag IP, and analyzed by mass-spectrometry as in Figure 1B. Illustrated residues on the domain-map of Stu2 indicate phosphorylated Serine and Threonine residues identified by mass-spectrometry. C. Cell viability of *STU2* mutants as in Fig. 1C, but with more conditions. Wild-type (M3), *stu2-AID* (no covering allele, M619), and *stu2-AID* cells expressing various *STU2-3HA* alleles from an _ectopic locus (*STU2*_*^WT^*_, M2898; *stu2*_*T836E S839D S840D S842D S852D S855D S858D,* _M3352*; stu2*_*T866E S867D T868E S880D T885E T886E*_, M3353; *stu2*_*T866E S867D T868E*_, M3354; *stu2*_*S880D T885E T886E T888E*_, M3355; *stu2*_*T866E S867D,* M3356; *stu2^S867D T868E^,* M3357; *stu2^T866E,T868E^,* M3358; *stu2^T866E^,* M2829) were serially diluted (five-fold) and spotted on plates containing DMSO, 500 μM auxin, 5 μg/mL benomyl, or 500 μM auxin + 5 μg/mL benomyl. D. Schematic representation of quantitative chromosome transmission fidelity assay adapted from (Zhu *et al*., 2015).

**Figure S2.**
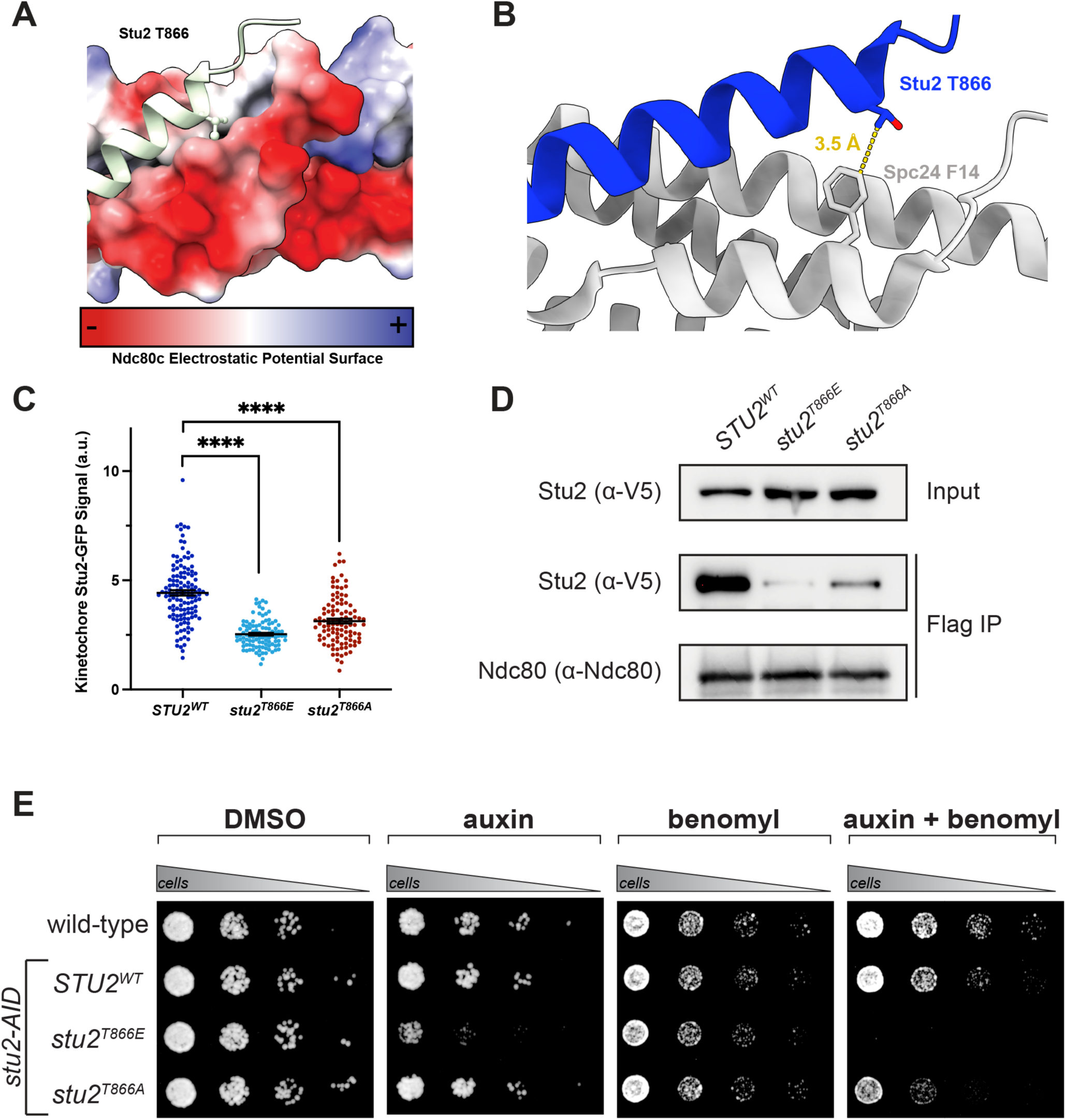
Stu2:Ndc80c binding details and *stu2^T866A^* mutant phenotypes. A. The crystal structure of Stu2 C-terminus bound to Ndc80^Dwarf^ (PDB 7KDF) showing electrostatic potential surface of Ndc80^Dwarf^. Stu2 illustrated as ribbon. B. Zoom in of crystal structure (PDB 7KDF) showing hydrophobic interaction between gamma carbon of Stu2^T866^ and Spc24^F14^. This carbon is lost in a *stu2^T866A^* mutation but preserved in *_stu2_T866V*. C. Exponentially growing *stu2-AID pMET-CDC20* cultures with an ectopically expressed *STU2-GFP* allele (*STU2^WT^-GFP*, M2599; *stu2^T866E^-GFP*, M2600, *stu2^T866A^-GFP,* M2601) that also contained *SPC110-mCherry* (spindle pole) were cultured in methionine-and auxin-containing media to arrest cells in metaphase. Cells were fixed and imaged to determine Kinetochore-Proximal Stu2-GFP. Bars represent mean of n=102-126 individual measurements. Error bars are S.E.M. p-values from two-tailed unpaired t-tests (*STU2^WT^* vs *stu2^T866E^*, p<0.0001; *STU2^WT^* vs *stu2^T866A^*, p<0.0001). D. Exponentially growing *stu2-AID* cultures expressing an ectopic copy of *STU2-3V5* (*STU2^WT^*, M622; *stu2^T866E^*, M1448; *stu2^T866A^*, M2108) as well as *DSN1-6His-3Flag* from the genomic locus were treated with auxin 30 min prior to harvesting. Kinetochore particles were purified from lysates by anti-Flag immunoprecipitation (IP) and analyzed by immunoblotting. E. Wild-type (M3), and *stu2-AID* cells expressing various *STU2-3HA* alleles from an ectopic locus (*STU2^WT^*, M2898; *stu2^T866E^,* M2829; *stu2^T866A^,* M2830) were serially diluted (fivefold) and spotted on plates containing DMSO, 500 μM auxin, 5 μg/mL benomyl, and 500 μM auxin + 5 μg/mL benomyl.

**Figure S3.**
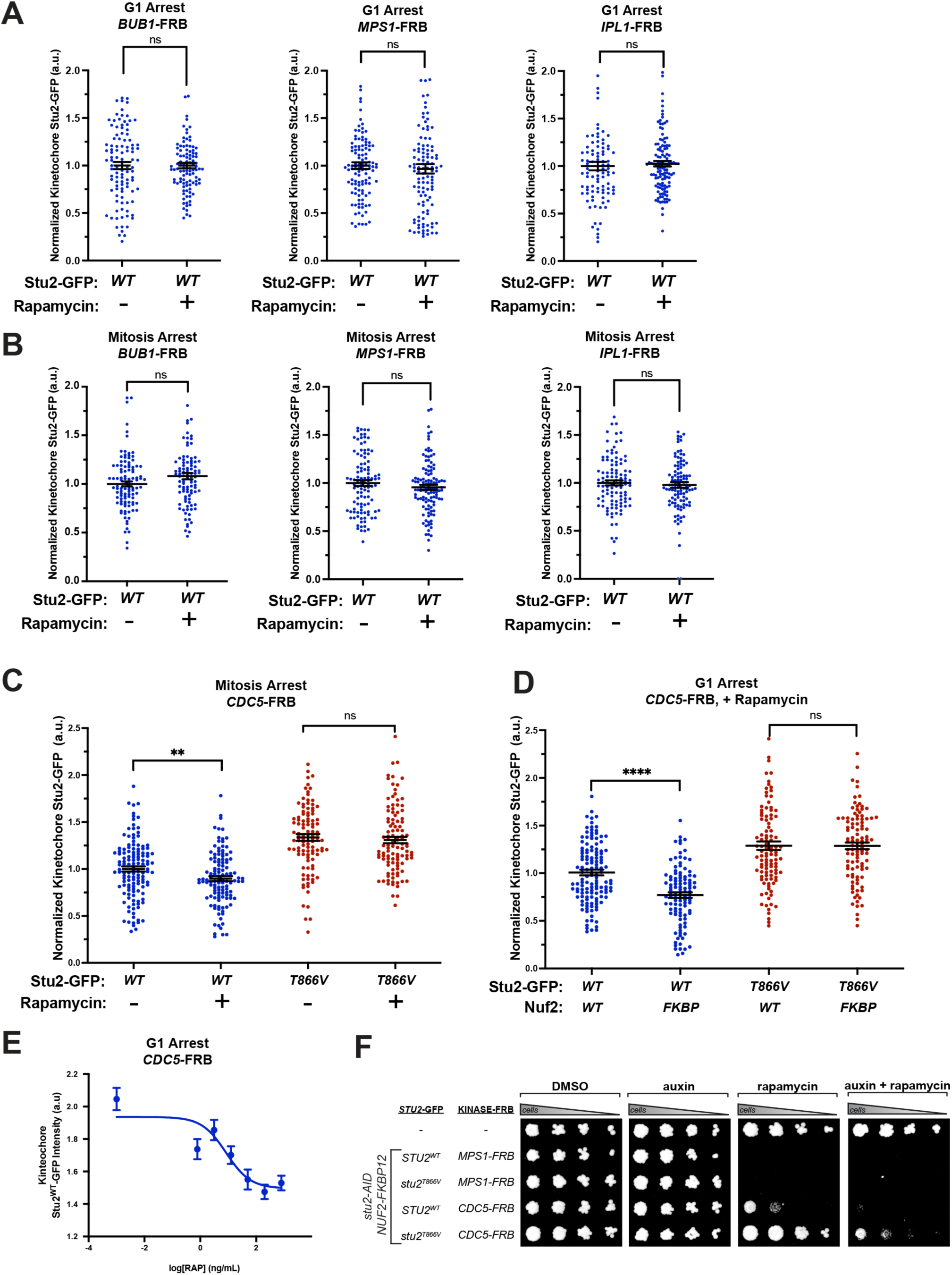
Effects of tethering mitotic kinases on Stu2 kinetochore association. A. Exponentially growing *stu2-AID* cells harboring *TOR1-1 fpr1Δ NUF2-FKBP12,* ectopic *STU2-GFP,* and a kinase-FRB allele (*BUB1-FRB*, M4970; *IPL1-FRB,* M4972*; MPS1-FRB*, M4792) were arrested in alpha factor for 3 hours. Cells received either 500 µM auxin + 200 ng/mL rapamycin or 500 µM auxin + DMSO for 30 minutes prior to being fixed and imaged. Bar represents mean of n=94-123 individual measurements. Error bars are S.E.M. p-values from two-tailed unpaired t-tests. (*BUB1-FRB* DMSO vs *BUB1-FRB* RAP, p=0.9961; *MPS1-FRB* DMSO vs *MPS1-FRB* RAP, p=0.6295; *IPL1-FRB* DMSO vs *IPL1-FRB* RAP, p=0.5822) B. Strains as in A were arrested in nocodazole for 2.5 hours. Cells received either 500 µM auxin + 200 ng/mL rapamycin or 500 µM auxin + DMSO for 30 minutes prior to being fixed and imaged. Bar represents average of n=99-113 individual measurements. Error bars are S.E.M. p-values from two-tailed unpaired t-tests. (*BUB1-FRB* DMSO vs *BUB1-FRB* RAP, p=0.0508; *MPS1-FRB* DMSO vs *MPS1-FRB* RAP, p=0.2715; *IPL1-FRB* DMSO vs *IPL1-FRB* RAP, p=0.5942) C. Exponentially growing *stu2-AID* cells harboring *TOR1-1 fpr1Δ NUF2-FKBP12 CDC5-FRB,* and an ectopic *STU2-GFP* variant (*STU2^WT^-GFP,* M4968; *stu2^T866V^-GFP,* M4969) were arrested in nocodazole for 2.5 hours. Cells received either 500 µM auxin + 200 ng/mL rapamycin or 500 µM auxin + DMSO for 30 minutes prior to being fixed and imaged. Bar represents mean of n=108-131 individual measurements. Error bars are S.E.M. p-values from two-tailed unpaired t-tests (*STU2^WT^* DMSO vs *STU2^WT^* RAP, p=0.0073; *stu2^T866V^* DMSO vs *stu2^T866V^* RAP, p=0.5791). D. Exponentially growing *stu2-AID* cells harboring *TOR1-1 fpr1Δ CDC5-FRB*) were arrested in alpha factor for 3 hours. Cells also contained *Stu2^WT^-GFP* with *NUF2^WT^* (M5094) or *NUF2-FKBP12* (M4968) or ectopic *stu2^T866V^-GFP* with *NUF2^WT^* (M5095) or *NUF2-FKBP12* (M4969) and received either 500 µM auxin + 200 ng/mL rapamycin or 500 µM auxin + DMSO for 30 minutes prior to being fixed and imaged. Bar represents mean of n=104-132 individual measurements. Error bars are S.E.M. p-values from two-tailed unpaired t-tests. (*STU2^WT^ NUF2^WT^* vs *STU2^WT^ NUF2-FKBP12*, p<0.0001; *stu2^T866V^ NUF2^WT^* vs *stu2^T866V^ NUF2-FKBP12*, p=0.9851). E. Exponentially growing *stu2-AID* cells harboring *TOR1-1 fpr1Δ NUF2-FKBP12 CDC5-FRB,* and ectopic *STU2^WT^-GFP* (M4968) were arrested in alpha factor for 3 hours. Cells received 500 µM auxin + varying dose of rapamycin (800 ng/mL, 200 ng/mL, 50 ng/mL, 12.5 ng/mL, 3.12 ng/mL, 0.78 ng/mL, 0 ng/mL) 30 minutes prior to being fixed and imaged. Points represent mean of n=120-208 individual measurements. Error bars are S.E.M. F. Cells harboring *TOR1-1 and fpr1Δ (*M1375), with *NUF2-FKBP12 MPS1-FRB stu2-AID* and *STU2^WT^-GFP* (M4792) or *stu2^T866V^-GFP* (M4793), or with *NUF2-FKBP12 CDC5-FRB stu2-AID* and *STU2^WT^-GFP* (M4968) or *stu2^T866V^-GFP* (M4969) were serially diluted and spotted on plates containing DMSO, auxin, rapamycin, and auxin + rapamycin. Tethering Mps1 to kinetochores has been previously show to result in spindle assembly checkpoint dependent cell death (Aravamudhan, Goldfarb and Joglekar, 2015).

**Figure S4.**
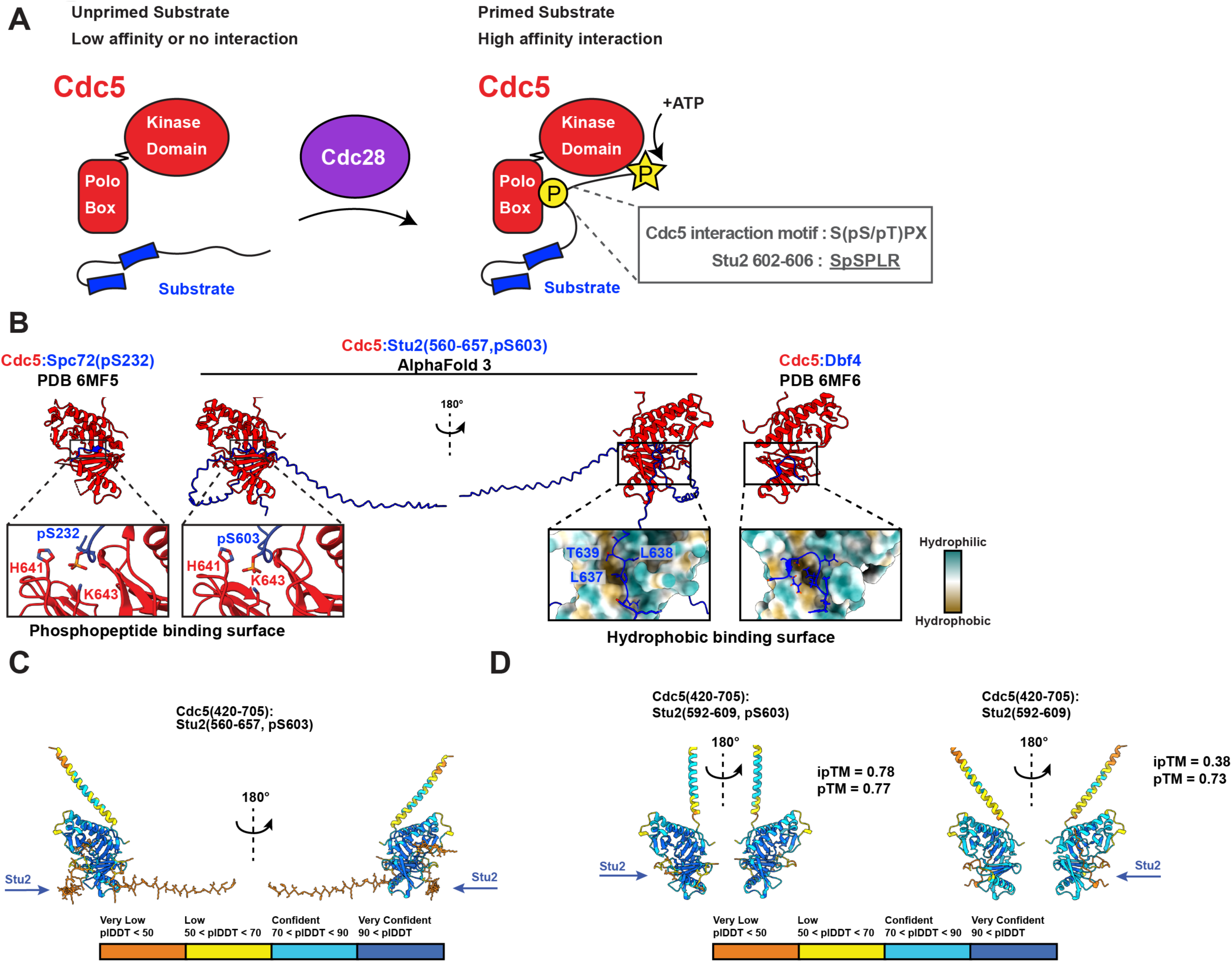
Different regions of Stu2 basic linker associate with Cdc5 polo box domain. A. Schematic representation of Cdk/Cdc28 priming substrates to interact with Cdc5. First, an unprimed substrate has low affinity or no interaction with Cdc5. Following Cdc28 phosphorylation (P in circle), substrates can interact with Cdc5 higher affinity, and Cdc5 adds its catalytic phosphorylation (P in star). B. LEFT – Crystal structure of Cdc5 polo-box domain bound to Spc72 (PDB 6MF5) (Almawi *et al*., 2020). Zoom in shows phosphorylated serine bound by lysine and histidine in Cdc5. MIDDLE – AlphaFold 3 prediction of Stu2 basic linker with pS603 (Stu2^560-657,pS603^) bound to Cdc5 polo box domain, as in Fig. 4B. Left shows pS603 binding to phosphopeptide binding residues on Cdc5 as in 6MF5. Right shows hydrophobic residues interacting with a hydrophobic patch on Cdc5. RIGHT – Crystal structure of Cdc5 polo-box domain bound to Dbf4 (PDB 6MF6) (Almawi *et al*., 2020). Zoom in shows Dbf4 interacting with hydrophobic surface on Cdc5. Stu2^600-607^ shows good agreement with the binding mode of Spc72 (LEFT) and Stu2^633-647^ interacts similar to the binding mode of Dbf4 (RIGHT) C. AlphaFold 3 prediction of Stu2 basic linker with pS603 (Stu2^560-657,pS603^) bound to Cdc5 polo box domain. Left shows pS603 binding to phosphopeptide residues on Cdc5 as in 6MF5. Right shows hydrophobic residues interacting with a hydrophobic patch on Cdc5. Colored by plDDT confidence score. Arrows indicate which surfaces Stu2 binds of Cdc5. D. LEFT: AlphaFold 3 prediction of Stu2 basic linker patch with pS603 (Stu2^592-607,pS603^) bound to Cdc5 polo box domain. Arrow indicates which surfaces Stu2 binds of Cdc5. RIGHT: AlphaFold 3 prediction of Stu2 basic linker patch without pS603 (Stu2^592-607^) bound to Cdc5 polo box domain. Arrow indicates which surfaces Stu2 binds of Cdc5. iPTM and PTM binding scores shown. Structures colored by plDDT confidence score.

**Figure S5.**
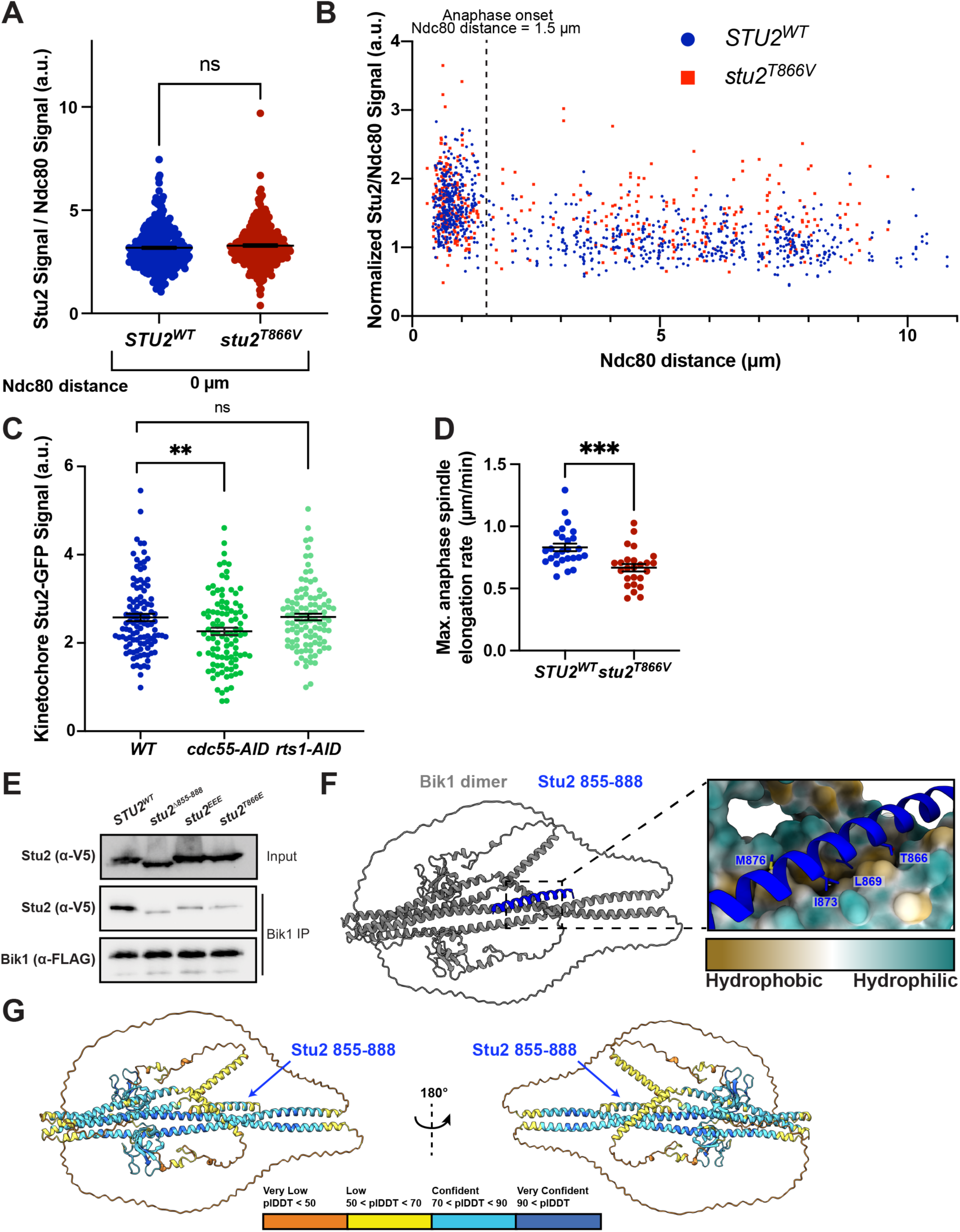
Cell cycle timing of Stu2^T866^ modification, PP2A regulatory subunit activity on Stu2, and predicted Stu2:Bik1 interaction. A. Exponentially growing *stu2-AID* cells expressing *STU2-GFP and NDC80-mKate* (M3774) or *stu2^T866V^-GFP* and *NDC80-mKate* (M4429), were released from a G1 arrest into auxin-containing media. Samples were taken every 15 minutes, fixed, and imaged. Ratio of Stu2/Ndc80c signal plotted for cells without separated/bilobed Ndc80 signal. Bars represent mean of n=274-337 individual measurements. Error bars are S.E.M. p-values from using a two-tailed unpaired t-test (*STU2^WT^* vs *stu2^T866V^*, p=0.1993). B. Data as in Fig. 6A but without binning by Ndc80c distance. Exponentially growing *stu2-AID* cells expressing *STU2-GFP and NDC80-mKate* (M3774) or *stu2^T866V^-GFP* and *NDC80-mKate* (M4429), were released from a G1 arrest into auxin-containing media. Samples were taken every 15 minutes, fixed, and imaged. Ratio of Stu2/Ndc80c signal plotted against the distance between corresponding Ndc80 puncta for cells with bilobed Ndc80 signal. C. Exponentially growing *stu2-AID* cells expressing ectopic *STU2-GFP* (M5505) or with *cdc55-AID* (M5527) or *rts1-AID* (M5506) were treated with nocodazole for 2.5 hours to arrest in mitosis. Cells treated with auxin 30 minutes prior to harvesting and fixed and imaged. Kinetochore associated Stu2-GFP signal was quantified. Bars represent mean from n=100-106 individual measurements. p-values are from an unpaired two-tailed t-test (*WT* vs *cdc55-AID*, p=0.0065; *WT* vs *rts1-AID,* p=0.0173). D. Cells grown and imaged in Fig. 7E were analyzed to determine maximum rates of spindle elongation over a 2-minute period for each individual cell. Each data point represents a single cell. Bars represent the average of n=25-26 individual measurements. Error bars are S.E.M. p-values from two-tailed unpaired t-test (*STU2^WT^* vs *stu2^T866V^*, p=0.0004). E. Exponentially growing *stu2-AID BIK1-3FLAG* cells expressing *STU2-3V5* variants (*STU2^WT^*, M1035; *stu2^Δ855-888^*, M1037; *stu2^L869E I873E M876E^*, M2285; *stu2^T866E^*, M2286) were treated with auxin 30 min prior to harvesting. Bik1 complexes were purified from lysates by anti-Flag immunoprecipitation (IP) and analyzed by immunoblotting. The loss of Stu2:Bik1 interaction in *stu2^T866E^* mutant cells implies that phosphorylation of Stu2^T866^ may alter interaction with Bik1, and potentially other Stu2 interactors that bind to Stu2’s C-terminus. F. AlphaFold 3 prediction of full-length Bik1 dimer bound to Stu2^(855-888)^. Zoom in shows hydrophobic residues on Stu2 interacting with hydrophobic Bik1 surface. G. Structure prediction as in F but colored to show plDDT confidence score.

**Table S1:** Yeast Strains used in this study.

| Strain | Relevant Genotype | Figure |
| --- | --- | --- |
| M3<br>(W303) | *MATa ura3-1 leu2-3,112 his3-11 trp1-1 can1-100 ade2-1 bar1-1* | 1C, S1C, S2E |
| M498 | *MATa STU2-3FLAG:KanMX* | 1B, S1B |
| M619 | *MATa his3::pGPD1-OsTIR1:HIS3 STU2-3HA-IAA7:KanMx DSN1-HIS-FLAG:URA3* | 1C, S1C |
| M622 | *MATa his3::pGPD1-OsTIR1:HIS3 STU2-3HA-IAA7:KanMx DSN1-HIS-FLAG:URA3 leu2::pSTU2-STU2-3V5:LEU2* | 2E, S2D |
| M1035 | *MATa his3::pGPD1-OsTIR1:HIS3 STU2-3HA-IAA7:KanMx BIK1-3FLAG:TRP1 leu2::pSTU2-STU2-3V5:LEU2* | S6A |
| M1037 | *MATa his3::pGPD1-OsTIR1:HIS3 STU2-3HA-IAA7:KanMx BIK1-3FLAG:TRP1 leu2::pSTU2-stu2(Δ855-888)-3V5:LEU2* | S6A |
| M1174 | *MATa SPC110-mCherry:HygMx MTW1-3GFP:HIS3* | 7F, 7G |
| M1375 | *MATa TOR1-1 fpr1Δ::NatMx* | 3D, S3F |
| M1448 | *MATa his3::pGPD1-OsTIR1:HIS3 STU2-3HA-IAA7:KanMx DSN1-HIS-FLAG:URA3 leu2::pSTU2-stu2(T866E)-3V5:LEU2* | 2E, S2D |
| M2108 | *MATa his3::pGPD1-OsTIR1:HIS3 STU2-3HA-IAA7:KanMx DSN1-HIS-FLAG:URA3 leu2::pSTU2-stu2(T866A)-3V5:LEU2* | S2D |
| M2285 | *MATa his3::pGPD1-OsTIR1:HIS3 STU2-3HA-IAA7:KanMx BIK1-3FLAG:TRP1 leu2::pSTU2-stu2(L869E I873E M876E)-3V5:LEU2* | S6A |
| M2286 | *MATa his3::pGPD1-OsTIR1:HIS3 STU2-3HA-IAA7:KanMx BIK1-3FLAG:TRP1 leu2::pSTU2-stu2(T866E)-3V5:LEU2* | S6A |
| M2298 | *MATa trp1::pGPD1-OsTIR1:TRP1 STU2-3HA-IAA7:KanMx leu2::pSTU2-STU2-GFP:LEU2 NUP2-mKate2:HisMx6* | 5D |
| M2351 | *MATa trp1::pGPD1-OsTIR1:TRP1 STU2-3HA-IAA7:KanMx leu2::pSTU2-stu2(S603A)-GFP:LEU2 NUP2-mKate2:HisMx6* | 5D |
| M2390 | *MATa trp1::pGPD1-OsTIR1:TRP1 STU2-3HA-IAA7:KanMx CDC20-AID:KanMx leu2::pSTU2-GFP-GST:LEU2 NUP2-mKate2:HisMx6* | 5A, 5B |
| M2391 | *MATa trp1::pGPD1-OsTIR1:TRP1 STU2-3HA-IAA7:KanMx CDC20-AID:KanMx leu2::pSTU2-NLS-GFP-GST:LEU2 NUP2-mKate2:HisMx6* | 5B |
| M2392 | *MATa trp1::pGPD1-OsTIR1:TRP1 STU2-3HA-IAA7:KanMx CDC20-AID:KanMx leu2::pSTU2-stu2(592-607)-GFP-GST:LEU2 NUP2-mKate2:HisMx6* | 5A, 5B, 5C |
| M2429 | *MATa trp1::pGPD1-OsTIR1:TRP1 STU2-3HA-IAA7:KanMx SPC110-mCherry:HygMx leu2::pSTU2-STU2-GFP:LEU2* | 7D, S5D |
| M2437 | *MATa trp1::pGPD1-OsTIR1:TRP1 STU2-3HA-IAA7:KanMx CDC20-AID:KanMx leu2::pSTU2-stu2(592-607 K598A R599A)-GFP-GST:LEU2 NUP2-mKate2:HisMx6* | 5B |
| M2438 | MATa *trp1::pGPD1-OsTIR1:TRP1 STU2-3HA-IAA7:KanMx CDC20-AID:KanMx leu2::pSTU2-stu2(592-607 S603A)-GFP-GST:LEU2 NUP2-mKate2:HisMx6* | 5C |
| M2439 | MATa *trp1::pGPD1-OsTIR1:TRP1 STU2-3HA-IAA7:KanMx leu2::pSTU2-stu2(592-607 K598A R599A)-GFP-GST:LEU2 NUP2-mKate2:HisMx6* | 5B |
| M2441 | MATa *trp1::pGPD1-OsTIR1:TRP1 STU2-3HA-IAA7:KanMx leu2::pSTU2-GFP-GST:LEU2 NUP2-mKate2:HisMx6* | 5B |
| M2442 | MATa *trp1::pGPD1-OsTIR1:TRP1 STU2-3HA-IAA7:KanMx leu2::pSTU2-NLS-GFP-GST:LEU2 NUP2-mKate2:HisMx6* | 5B |
| M2443 | MATa *trp1::pGPD1-OsTIR1:TRP1 STU2-3HA-IAA7:KanMx leu2::pSTU2-stu2(592-607)-GFP-GST:LEU2 NUP2-mKate2:HisMx6* | 5B |
| M2599 | MATa *his3::pGPD1-OsTIR1:HIS3 STU2-3HA-IAA7:KanMx SPC110-mCherry:HygMx pMET-CDC20:TRP1 leu2::pSTU2-STU2-GFP:LEU2* | 2F, S2C |
| M2600 | MATa *his3::pGPD1-OsTIR1:HIS3 STU2-3HA-IAA7:KanMx SPC110-mCherry:HygMx pMET-CDC20:TRP1 leu2::pSTU2-stu2(T866E)-GFP:LEU2* | 2F, S2C |
| M2601 | MATa *his3::pGPD1-OsTIR1:HIS3 STU2-3HA-IAA7:KanMx SPC110-mCherry:HygMx pMET-CDC20:TRP1 leu2::pSTU2-stu2(T866A)-GFP:LEU2* | S2C |
| M2726 | MATa *trp1::pGPD1-OsTIR1:TRP1 STU2-3HA-IAA7:KanMx CDC20-AID:KanMx leu2::pSTU2-stu2(592-607 S603E)-GFP-GST:LEU2 NUP2-mKate2:HisMx6* | 5C |
| M2827 | MATa *pMET-CDC20:TRP1 STU2-3GFP:His3Mx* | 3C |
| M2829 | MATa *his3::pGPD1-OsTIR1:HIS3 STU2-3HA-IAA7:KanMx DSN1-HIS-FLAG:URA3 trp1::pSTU2-stu2(T866E)-3HA:TRP1* | 1C, S1C, S2E |
| M2830 | MATa *his3::pGPD1-OsTIR1:HIS3 STU2-3HA-IAA7:KanMx DSN1-HIS-FLAG:URA3 trp1::pSTU2-stu2(T866A)-3HA:TRP1* | S2E |
| M2898 | MATa *his3::pGPD1-OsTIR1:HIS3 STU2-3HA-IAA7:KanMx DSN1-HIS-FLAG:URA3 trp1::pSTU2-STU2-3HA:TRP1* | 1C, S1C, S2E |
| M3138 | MATa *pMET-CDC20:TRP1 STU2-3GFP:His3Mx cdc5-1* | 3C |
| M3276 | MATa *his3::pGPD1-OsTIR1:HIS3 STU2-3V5-IAA7:KanMx MFA1-3XGFP:HIS5 MC(CEN3.L.YFS5.1)MATα:LEU2* | 1C, S1C |
| M3352 | MATa *his3::pGPD1-OsTIR1:HIS3 STU2-3HA-IAA7:KanMx DSN1-HIS-FLAG:URA3 trp1::pSTU2-stu2(T836E S839D S840D S842D S852D S855D S858D)-3HA:TRP1* | 1C, S1C |
| M3353 | MATa *his3::pGPD1-OsTIR1:HIS3 STU2-3HA-IAA7:KanMx DSN1-HIS-FLAG:URA3 trp1::pSTU2-stu2(T866E S867D T868E S880D T885E T886E)-3HA:TRP1* | 1C, S1C |
| M3354 | MATa *his3::pGPD1-OsTIR1:HIS3 STU2-3HA-IAA7:KanMx DSN1-HIS-FLAG:URA3 trp1::pSTU2-stu2(T866E S867D T868E)-3HA:TRP1* | 1C, S1C |
| M3355 | MATa *his3::pGPD1-OsTIR1:HIS3 STU2-3HA-IAA7:KanMx DSN1-HIS-FLAG:URA3 trp1::pSTU2-stu2(S880D T885E T886E T888E)-3HA:TRP1* | 1C, S1C |
| M3356 | MATa *his3::pGPD1-OsTIR1:HIS3 STU2-3HA-IAA7:KanMx DSN1-HIS-FLAG:URA3 trp1::pSTU2-stu2(T866E S867D)-3HA:TRP1* | 1C, S1C |
| M3357 | MATa his3::pGPD1-OsTIR1:HIS3 STU2-3HA-IAA7:KanMx DSN1-HIS-FLAG:URA3 trp1::pSTU2-stu2(S867D T868E)-3HA:TRP1 | 1C, S1C |
| M3358 | MATa his3::pGPD1-OsTIR1:HIS3 STU2-3HA-IAA7:KanMx DSN1-HIS-FLAG:URA3 trp1::pSTU2-stu2(T866E T868E)-3HA:TRP1 | 1C, S1C |
| M3451 | MATa his3::pGPD1-OsTIR1:HIS3 STU2-3V5-IAA7:KanMx MFA1-3XGFP:HIS5 MC(CEN3.L.YFS5.1)MATα:LEU2 trp1::pSTU2-STU2-3HA:TRP1 | 1C, S1C |
| M3452 | MATa his3::pGPD1-OsTIR1:HIS3 STU2-3V5-IAA7:KanMx MFA1-3XGFP:HIS5 MC(CEN3.L.YFS5.1)MATα:LEU2 trp1::pSTU2-stu2(T866E)-3HA:TRP1 | 1C, S1C |
| M3592 | MATa his3::pGPD1-OsTIR1:HIS3 STU2-3V5-IAA7:KanMx MFA1-3XGFP:HIS5 MC(CEN3.L.YFS5.1)MATα:LEU2 trp1::pSTU2-stu2(T836E S839D S840D S842D S852D S855D S858D)-3HA:TRP1 | 1C, S1C |
| M3593 | MATa his3::pGPD1-OsTIR1:HIS3 STU2-3V5-IAA7:KanMx MFA1-3XGFP:HIS5 MC(CEN3.L.YFS5.1)MATα:LEU2 trp1::pSTU2-stu2(T866E S867D T868E S880D T885E T886E)-3HA:TRP1 | 1C, S1C |
| M3594 | MATa his3::pGPD1-OsTIR1:HIS3 STU2-3V5-IAA7:KanMx MFA1-3XGFP:HIS5 MC(CEN3.L.YFS5.1)MATα:LEU2 trp1::pSTU2-stu2(T866E S867D T868E)-3HA:TRP1 | 1C, S1C |
| M3595 | MATa his3::pGPD1-OsTIR1:HIS3 STU2-3V5-IAA7:KanMx MFA1-3XGFP:HIS5 MC(CEN3.L.YFS5.1)MATα:LEU2 trp1::pSTU2-stu2(S880D T885E T886E T888E)-3HA:TRP1 | 1C, S1C |
| M3596 | MATa his3::pGPD1-OsTIR1:HIS3 STU2-3V5-IAA7:KanMx MFA1-3XGFP:HIS5 MC(CEN3.L.YFS5.1)MATα:LEU2 trp1::pSTU2-stu2(T866E S867D)-3HA:TRP1 | 1C, S1C |
| M3597 | MATa his3::pGPD1-OsTIR1:HIS3 STU2-3V5-IAA7:KanMx MFA1-3XGFP:HIS5 MC(CEN3.L.YFS5.1)MATα:LEU2 trp1::pSTU2-stu2(S867D T868E)-3HA:TRP1 | 1C, S1C |
| M3598 | MATa his3::pGPD1-OsTIR1:HIS3 STU2-3V5-IAA7:KanMx MFA1-3XGFP:HIS5 MC(CEN3.L.YFS5.1)MATα:LEU2 trp1::pSTU2-stu2(T866E T868E)-3HA:TRP1 | 1C, S1C |
| M3774 | MATa trp1::pGPD1-OsTIR1:TRP1 STU2-3HA-IAA7 NDC80-mKate2:HisMX3 leu2::pSTU2-STU2-GFP:LEU2 | 1A, 6A, 6B, 7A, 7B, 7C, S1A, S5A, S5B |
| M4398 | MATa his3::pGPD1-OsTIR1:HIS3 STU2-3HA-IAA7:KanMx DSN1-HIS-FLAG:URA3 leu2::pSTU2-stu2(T866V)-3V5:LEU2 | 2E |
| M4429 | MATa trp1::pGPD1-OsTIR1:TRP1 STU2-3HA-IAA7 NDC80-mKate2:HisMX3 leu2::pSTU2-stu2(T866V)-GFP:LEU2 | 6A, 6B, 7A, 7B, 7C, S5A, S5B |
| M4447 | MATa his3::pGPD1-OsTIR1:HIS3 STU2-3HA-IAA7:KanMx SPC110-mCherry:HygMx pMET-CDC20:TRP1 leu2::pSTU2-stu2(T866V)-GFP:LEU2 | 2F |
| M4792 | MATa trp1::pGPD1-OsTIR1:TRP1 TOR1-1 fpr1Δ::NatMx NUF2-FKBP12:HIS MPS1-FRB:KanMx STU2-3HA-IAA7:KanMx leu2::pSTU2-STU2-GFP:LEU2 DSN1-HIS-FLAG:URA3 | S3A, S3B, S3F |
| M4793 | *MATa trp1::pGPD1-OsTIR1:TRP1 TOR1-1 fpr1Δ::NatMx NUF2-FKBP12:HIS MPS1-FRB:KanMx STU2-3HA-IAA7:KanMx leu2::pSTU2-stu2(T866V)-GFP:LEU2 DSN1-HIS-FLAG:URA3* | S3F |
| M4968 | *MATa trp1::pGPD1-OsTIR1:TRP1 TOR1-1 fpr1Δ::NatMx NUF2-FKBP12:HIS CDC5-FRB:KanMx STU2-3HA-IAA7:KanMx leu2::pSTU2-STU2-GFP:LEU2* | 3B, 3D,<br>S3C,<br>S3D,<br>S3E, S3F |
| M4969 | *MATa trp1::pGPD1-OsTIR1:TRP1 TOR1-1 fpr1Δ::NatMx NUF2-FKBP12:HIS CDC5-FRB:KanMx STU2-3HA-IAA7:KanMx leu2::pSTU2-stu2(T866V)-GFP:LEU2* | 3B, 3D,<br>S3C,<br>S3D,<br>S3F |
| M4970 | *MATa trp1::pGPD1-OsTIR1:TRP1 TOR1-1 fpr1Δ::NatMx NUF2-FKBP12:HIS BUB1-FRB:KanMx STU2-3HA-IAA7:KanMx leu2::pSTU2-STU2-GFP:LEU2* | S3A,<br>S3B |
| M4972 | *MATa trp1::pGPD1-OsTIR1:TRP1 TOR1-1 fpr1Δ::NatMx NUF2-FKBP12:HIS IPL1-FRB:KanMx STU2-3HA-IAA7:KanMx leu2::pSTU2-STU2-GFP:LEU2* | S3A,<br>S3B |
| M5094 | *MATa trp1::pGPD1-OsTIR1:TRP1 TOR1-1 fpr1Δ::NatMx CDC5-FRB:KanMx STU2-3HA-IAA7:KanMx leu2::pSTU2-STU2-GFP:LEU2* | S3D |
| M5095 | *MATa trp1::pGPD1-OsTIR1:TRP1 TOR1-1 fpr1Δ::NatMx CDC5-FRB:KanMx STU2-3HA-IAA7:KanMx leu2::pSTU2-stu2(T866V)-GFP:LEU2* | S3D |
| M5145 | *MATa trp1::pGPD1-OsTIR1:TRP1 STU2-3V5-IAA7:KanMx SPC110-mCherry:HygMx CDC20-AID:KanMx leu2::pSTU2-STU2-NLS-GFP:LEU2 ura3-1::TUB1-CFP:URA3* | 4C |
| M5147 | *MATa trp1::pGPD1-OsTIR1:TRP1 STU2-3V5-IAA7:KanMx SPC110-mCherry:HygMx CDC20-AID:KanMx leu2::pSTU2-stu(S603A)-NLS-GFP:LEU2 ura3-1::TUB1-CFP:URA3* | 4C |
| M5149 | *MATa trp1::pGPD1-OsTIR1:TRP1 STU2-3V5-IAA7:KanMx SPC110-mCherry:HygMx CDC20-AID:KanMx leu2::pSTU2-stu(Δ600-605)-NLS-GFP:LEU2 ura3-1::TUB1-CFP:URA3* | 4C |
| M5150 | *MATa trp1::pGPD1-OsTIR1:TRP1 TOR1-1 fpr1Δ::NatMx NUF2-FKBP12:HIS CDC5-FRB:KanMx STU2-3HA-IAA7:KanMx leu2::pSTU2-STU2-NLS-GFP:LEU2* | 4E |
| M5154 | *MATa trp1::pGPD1-OsTIR1:TRP1 TOR1-1 fpr1Δ::NatMx NUF2-FKBP12:HIS CDC5-FRB:KanMx STU2-3HA-IAA7:KanMx leu2::pSTU2-stu2(Δ600-605)-NLS-GFP:LEU2* | 4E |
| M5309 | *MATa trp1::pGPD1-OsTIR1:TRP STU2-3HA-IAA7:KanMx SPC110-mCherry:HygMx leu2::pSTU2-stu2(T866V)-GFP:LEU2* | 7D, S5D |
| M5335 | *MATa trp1::pGPD1-OsTIR1:TRP1 STU2-3HA-IAA7:NatMx DSN1-HIS-FLAG:URA3 CDC20-AID:KanMx leu2::pSTU2-STU2-GFP:LEU2* | 6C |
| M5337 | *MATa trp1::pGPD1-OsTIR1:TRP1 STU2-3HA-IAA7:NatMx DSN1-HIS-FLAG:URA3 CDC20-AID:KanMx leu2::pSTU2-STU2-GFP:LEU2 cdc55Δ::KanMx* | 6C |
| M5505 | *MATa trp1::pGPD1-OsTIR1:TRP1 STU2-3HA-IAA7:NatMx leu2::pSTU2-STU2-GFP:LEU2* | S5C |
| M5506 | *MATa trp1::pGPD1-OsTIR1:TRP1 STU2-3HA-IAA7:NatMx leu2::pSTU2-STU2-GFP:LEU2 RTS1-3HA-IAA7:HygMx* | S5C |
| M5527 | *MATa trp1::pGPD1-OsTIR1:TRP1 STU2-3HA-IAA7:NatMx leu2::pSTU2-STU2-GFP:LEU2 CDC55-3HA-IAA7:HygMx* | S5C |
| M5554 | *MATa trp1::pGPD1-OsTIR1:TRP1 STU2-3HA-IAA7:NatMx DSN1-HIS-FLAG:URA3 leu2::pSTU2-STU2-NLS-GFP:LEU2* | 4D |
| M5555 | *MATa trp1::pGPD1-OsTIR1:TRP1 STU2-3HA-IAA7:NatMx DSN1-HIS-FLAG:URA3 leu2::pSTU2-STU2-NLS-GFP:LEU2 cdc28-as1* | 4D |
| M5611 | *MATa trp1::pGPD1-OsTIR1:TRP1 STU2-3HA-IAA7:NatMx DSN1-HIS-FLAG:URA3 CDC20-AID:KanMx leu2::pSTU2-stu2(T866V)-GFP:LEU2 cdc55Δ::KanMx* | 6C |
| M5762 | *MATa trp1::pGPD1-OsTIR1:TRP1 STU2-3HA-IAA7:KanMx leu2::pSTU2-STU2-GFP:LEU2 NUP2-mKate2:HisMx6* | 5E |
| M5770 | *MATa trp1::pGPD1-OsTIR1:TRP1 STU2-3HA-IAA7:KanMx leu2::pSTU2-STU2-GFP:LEU2 NUP2-mKate2:HisMx6 cdc28-as1* | 5E |
| M5848 | *MATa SPC110-mCherry:HygMx MTW1-3GFP:HIS3 cdc55Δ::KanMx* | 7F, 7G |
| M5980 | *MATa trp1::pGPD1-OsTIR1:TRP1 STU2-3V5-IAA7:KanMx SPC110-mCherry:HygMx CDC20-AID:KanMx leu2::pSTU2-stu(Δ633-647)-NLS-GFP:LEU2 ura3-1::TUB1-CFP:URA3* | 4C |
| M5989 | *MATa trp1::pGPD1-OsTIR1:TRP1 TOR1-1 fpr1Δ::NatMx NUF2-FKBP12:HIS CDC5-FRB:KanMx STU2-3HA-IAA7:KanMx leu2::pSTU2-stu2(Δ633-647)-NLS-GFP:LEU2* | 4E |

**Table S2:**
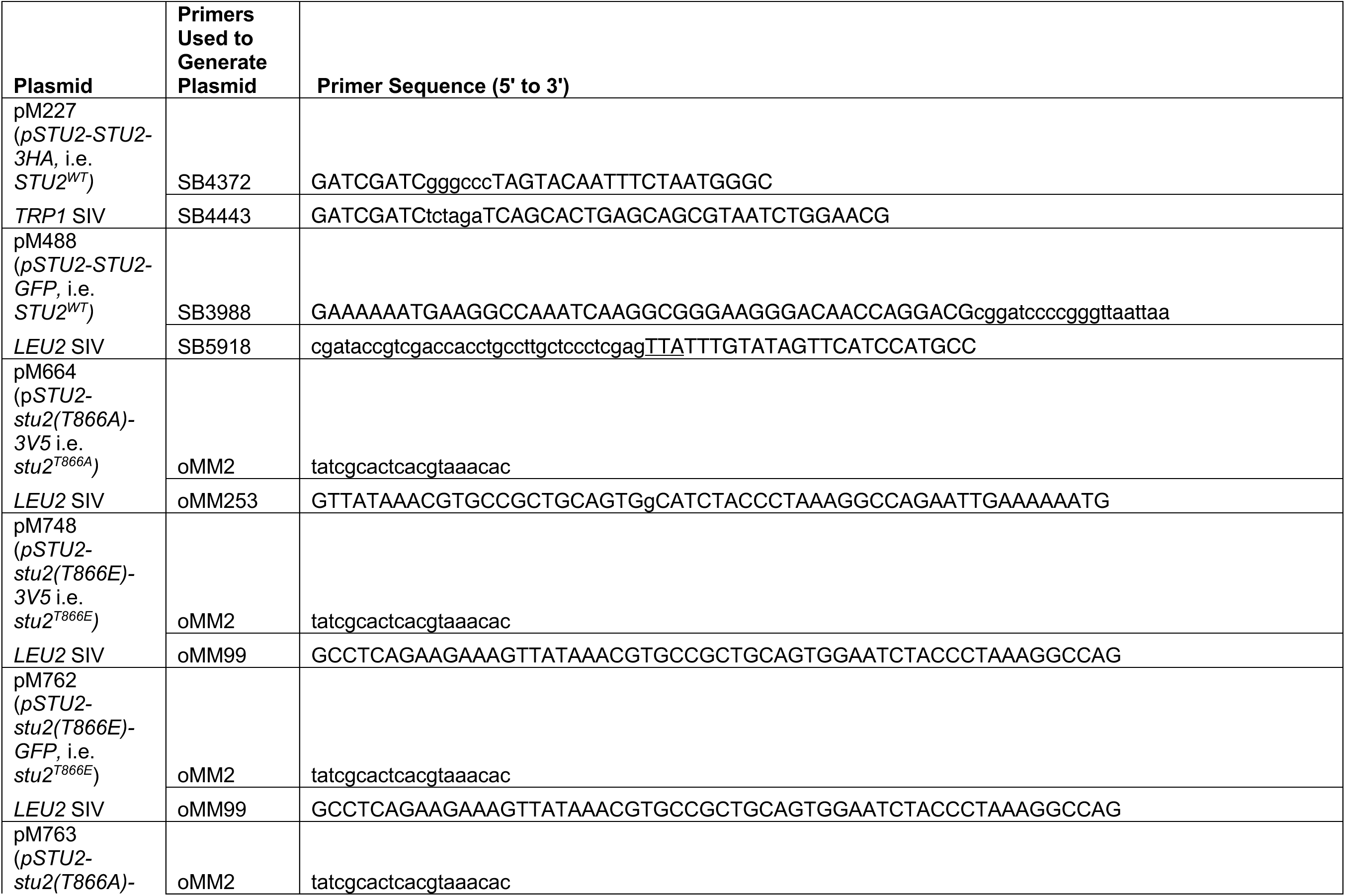
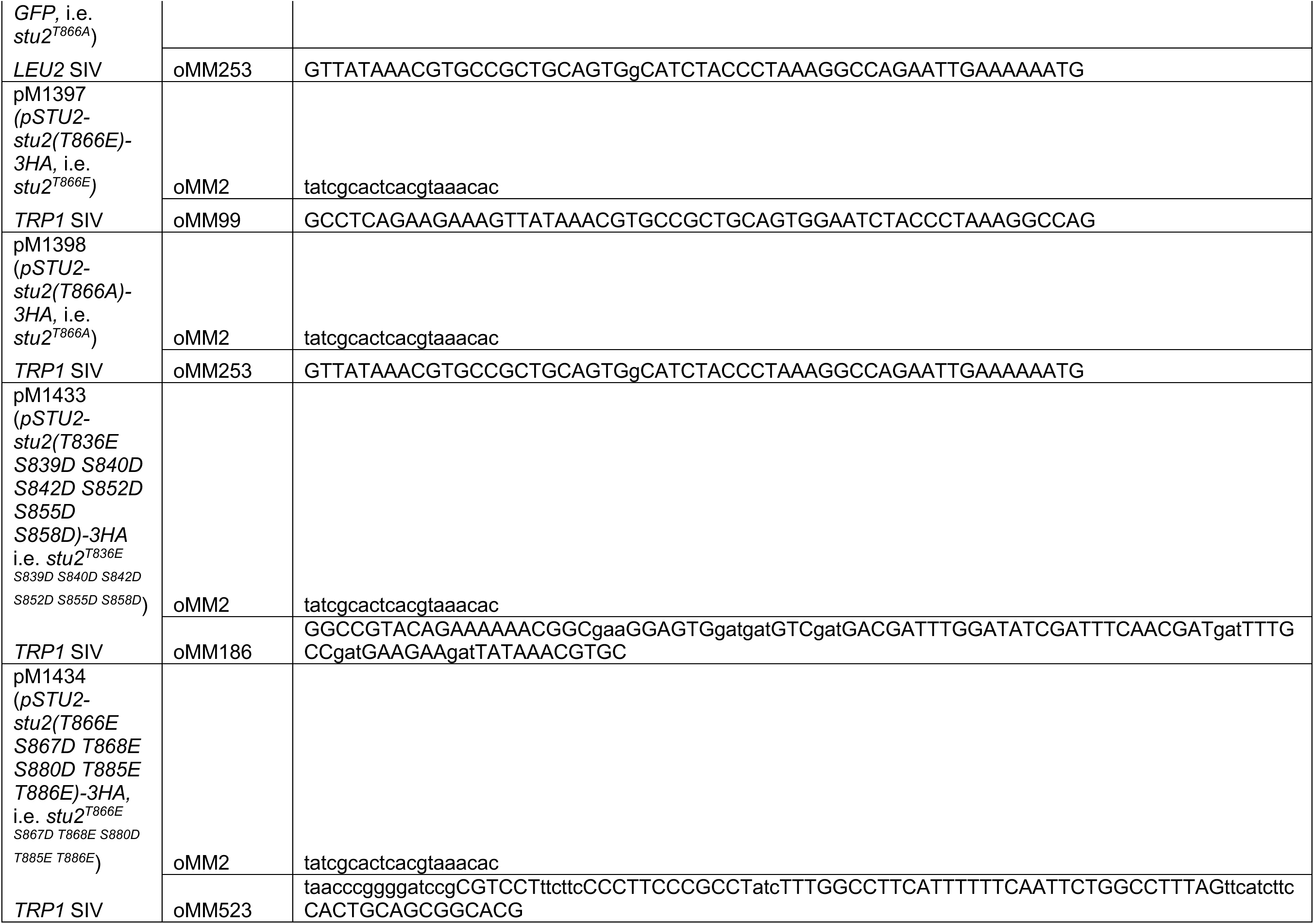
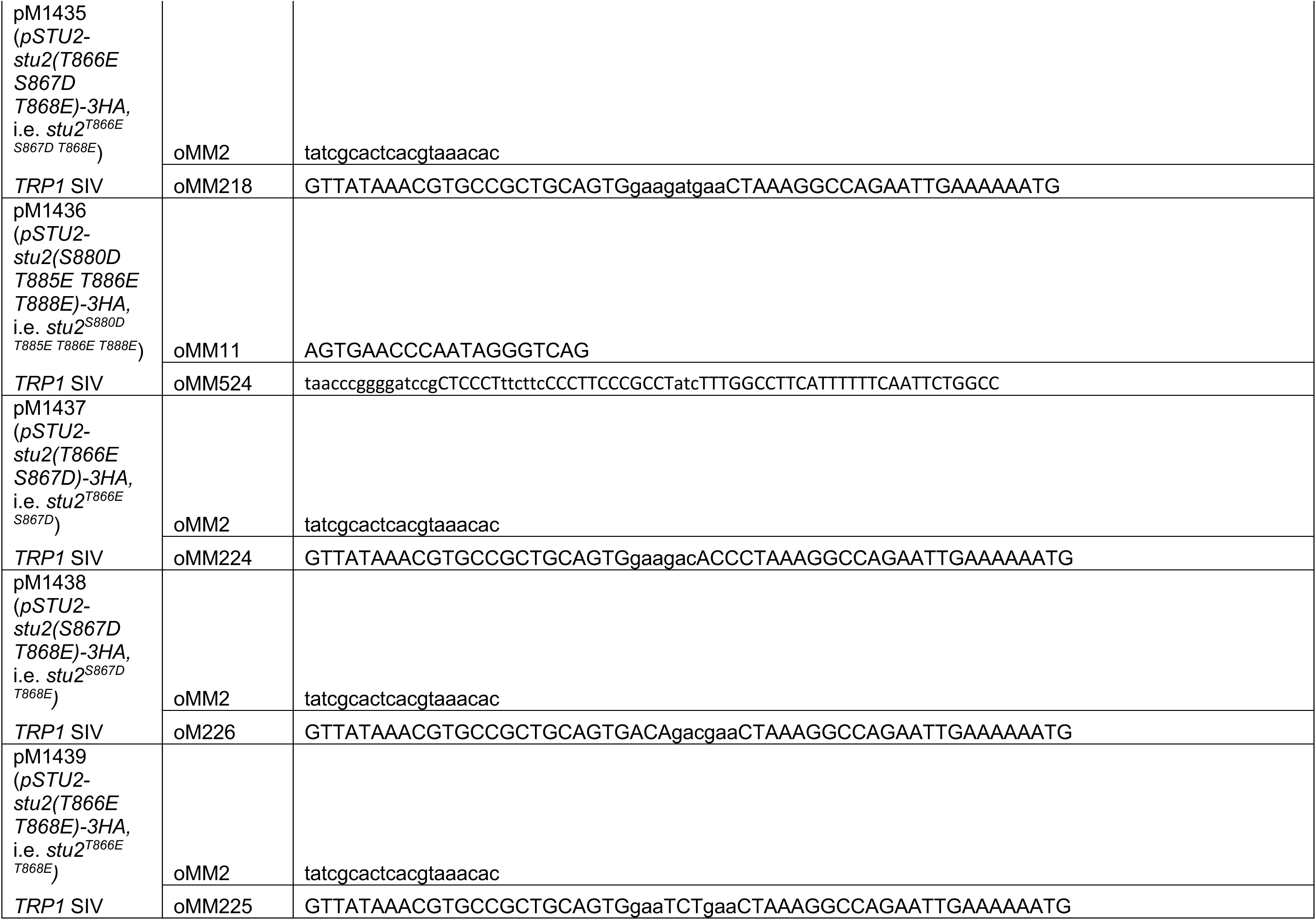
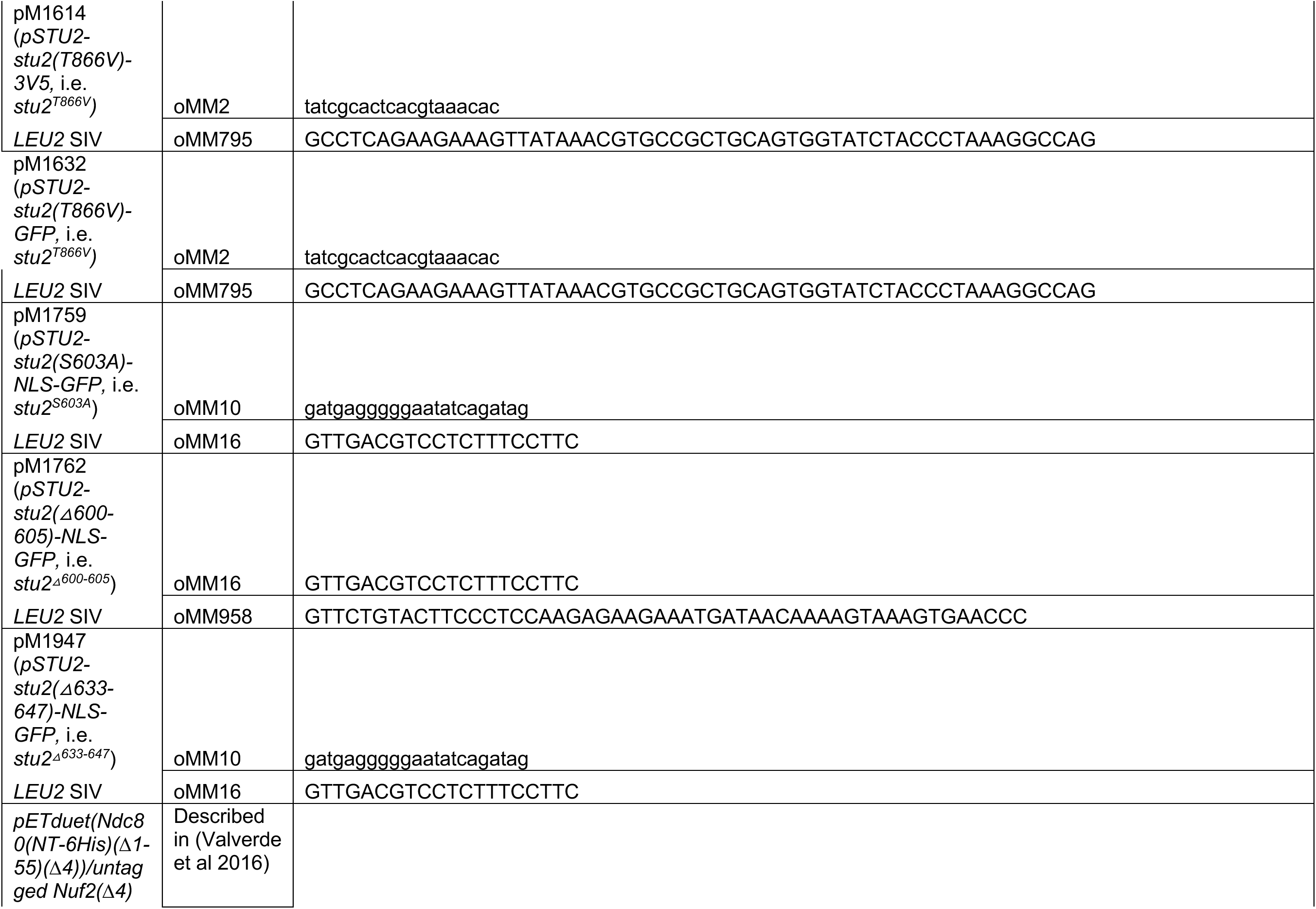
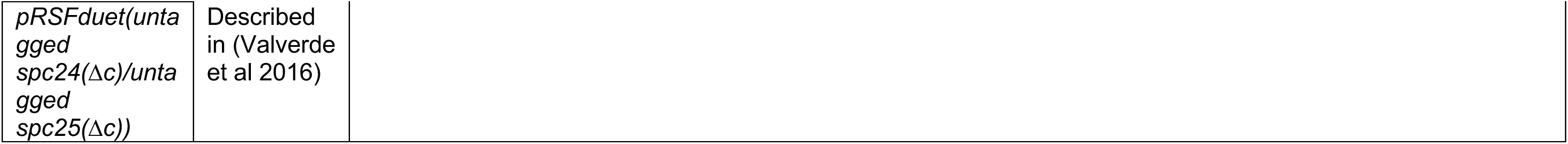
Plasmids and Oligos generated used this study.

## REFERENCES

Abramson, J. et al. (2024) ‘Accurate structure prediction of biomolecular interactions with AlphaFold 3’, Nature, 630(8016), pp. 493–500. Available at: 10.1038/s41586-024-07487-w.

Akiyoshi, B. et al. (2010) ‘Tension directly stabilizes reconstituted kinetochore-microtubule attachments’, Nature, 468(7323), pp. 576–579. Available at: 10.1038/nature09594.

Al-Bassam, J. et al. (2006) ‘Stu2p binds tubulin and undergoes an open-to-closed conformational change’, The Journal of Cell Biology, 172(7), pp. 1009–1022. Available at: 10.1083/jcb.200511010.

Almawi, A.W. et al. (2020) ‘Distinct surfaces on Cdc5/PLK Polo-box domain orchestrate combinatorial substrate recognition during cell division’, Scientific Reports, 10(1), p. 3379. Available at: 10.1038/s41598-020-60344-4.

Altschul, S. (1997) ‘Gapped BLAST and PSI-BLAST: a new generation of protein database search programs’, Nucleic Acids Research, 25(17), pp. 3389–3402. Available at: 10.1093/nar/25.17.3389.

Amin, M.A., Agarwal, S. and Varma, D. (2019) ‘Mapping the kinetochore MAP functions required for stabilizing microtubule attachments to chromosomes during metaphase’, Cytoskeleton, 76(6), pp. 398–412. Available at: 10.1002/cm.21559.

Aoki, K. et al. (2006) ‘Cdc2 phosphorylation of the fission yeast Dis1 ensures accurate chromosome segregation’, Current biology: CB, 16(16), pp. 1627–1635. Available at: 10.1016/j.cub.2006.06.065.

Aravamudhan, P. et al. (2014) ‘Assembling the protein architecture of the budding yeast kinetochore-microtubule attachment using FRET’, Current biology: CB, 24(13), pp. 1437–1446. Available at: 10.1016/j.cub.2014.05.014.

Aravamudhan, P., Goldfarb, A.A. and Joglekar, A.P. (2015) ‘The kinetochore encodes a mechanical switch to disrupt spindle assembly checkpoint signalling’, Nature Cell Biology, 17(7), pp. 868–879. Available at: 10.1038/ncb3179.

Ashburner, M. et al. (2000) ‘Gene Ontology: tool for the unification of biology’, Nature Genetics, 25(1), pp. 25–29. Available at: 10.1038/75556.

Ayaz, P. et al. (2012) ‘A TOG:αβ-tubulin complex structure reveals conformation-based mechanisms for a microtubule polymerase’, *Science (New York*, N.Y.), 337(6096), pp. 857–860. Available at: 10.1126/science.1221698.

Ayaz, P. et al. (2014) ‘A tethered delivery mechanism explains the catalytic action of a microtubule polymerase’, eLife, 3, p. e03069. Available at: 10.7554/eLife.03069.

Baro, B. et al. (2018) ‘SILAC-based phosphoproteomics reveals new PP2A-Cdc55-regulated processes in budding yeast’, GigaScience, 7(5). Available at: 10.1093/gigascience/giy047.

Bloom, J. and Cross, F.R. (2007) ‘Multiple levels of cyclin specificity in cell-cycle control’, Nature Reviews Molecular Cell Biology, 8(2), pp. 149–160. Available at: 10.1038/nrm2105.

van Breugel, M., Drechsel, D. and Hyman, A. (2003) ‘Stu2p, the budding yeast member of the conserved Dis1/XMAP215 family of microtubule-associated proteins is a plus end-binding microtubule destabilizer’, The Journal of cell biology, 161(2), pp. 359–369. Available at: 10.1083/jcb.200211097.

Burnette, W.N. (1981) ‘“Western blotting”: electrophoretic transfer of proteins from sodium dodecyl sulfate--polyacrylamide gels to unmodified nitrocellulose and radiographic detection with antibody and radioiodinated protein A’, Analytical Biochemistry, 112(2), pp. 195–203.

Buvelot, S. et al. (2003) ‘The budding yeast Ipl1/Aurora protein kinase regulates mitotic spindle disassembly’, The Journal of Cell Biology, 160(3), pp. 329–339. Available at: 10.1083/jcb.200209018.

Cairo, G., et al. (2023) ‘Distinct Aurora B pools at the inner centromere and kinetochore have different contributions to meiotic and mitotic chromosome segregation’, Molecular Biology of the Cell. Edited by K. Bloom, 34(5), p. ar43. Available at: 10.1091/mbc.E23-01-0014.

Calabria, I. et al. (2012) ‘Zds1 regulates PP2ACdc55 activity and Cdc14 activation during mitotic exit via its Zds_C motif’, *Journal of Cell Science*, p. jcs.097865. Available at: 10.1242/jcs.097865.

Carrier, J.S. et al. (2022) ‘Stimulating microtubule growth is not the essential function of the microtubule polymerase Stu2’. Available at: 10.1101/2022.09.09.507218.

Ciferri, C. et al. (2008) ‘Implications for kinetochore-microtubule attachment from the structure of an engineered Ndc80 complex’, Cell, 133(3), pp. 427–439. Available at: 10.1016/j.cell.2008.03.020.

Clift, D., Bizzari, F. and Marston, A.L. (2009) ‘Shugoshin prevents cohesin cleavage by PP2A ^Cdc55^ - dependent inhibition of separase’, Genes & Development, 23(6), pp. 766–780. Available at: 10.1101/gad.507509.

Conti, D. et al. (2024) ‘Role of PLK1 in the epigenetic maintenance of centromeres’. Available at: 10.1101/2024.02.23.581696.

De Gramont, A. and Cohen-Fix, O. (2005) ‘The many phases of anaphase’, Trends in Biochemical Sciences, 30(10), pp. 559–568. Available at: 10.1016/j.tibs.2005.08.008.

Dudziak, A. et al. (2021) ‘Phospho-regulated Bim1/EB1 interactions trigger Dam1c ring assembly at the budding yeast outer kinetochore’, The EMBO Journal, 40(18), p. e108004. Available at: 10.15252/embj.2021108004.

Elia, A.E.H. et al. (2003) ‘The Molecular Basis for Phosphodependent Substrate Targeting and Regulation of Plks by the Polo-Box Domain’, Cell, 115(1), pp. 83–95. Available at: 10.1016/S0092-8674(03)00725-6.

Fees, C.P., Estrem, C. and Moore, J.K. (2017) ‘High-resolution Imaging and Analysis of Individual Astral Microtubule Dynamics in Budding Yeast’, Journal of Visualized Experiments, (122), p. 55610. Available at: 10.3791/55610-v.

Goshima, G. and Vale, R.D. (2003) ‘The roles of microtubule-based motor proteins in mitosis’, The Journal of Cell Biology, 162(6), pp. 1003–1016. Available at: 10.1083/jcb.200303022.

Greenlee, M.A., et al. (2022) ‘The TOG protein Stu2 is regulated by acetylation’, PLOS Genetics. Edited by G.P. Copenhaver, 18(9), p. e1010358. Available at: 10.1371/journal.pgen.1010358.

Gunzelmann, J. et al. (2018) ‘The microtubule polymerase Stu2 promotes oligomerization of the γ-TuSC for cytoplasmic microtubule nucleation’, eLife, 7. Available at: 10.7554/eLife.39932.

Guo, Y. et al. (2006) ‘Analysis of cellular responses to aflatoxin B1 in yeast expressing human cytochrome P450 1A2 using cDNA microarrays’, Mutation Research/Fundamental and Molecular Mechanisms of Mutagenesis, 593(1–2), pp. 121–142. Available at: 10.1016/j.mrfmmm.2005.07.001.

Gutierrez, A. et al. (2020) ‘Cdk1 Phosphorylation of the Dam1 Complex Strengthens Kinetochore-Microtubule Attachments’, Current biology: CB [Preprint]. Available at: 10.1016/j.cub.2020.08.054.

Haruki, H., Nishikawa, J. and Laemmli, U.K. (2008) ‘The Anchor-Away Technique: Rapid, Conditional Establishment of Yeast Mutant Phenotypes’, Molecular Cell, 31(6), pp. 925–932. Available at: 10.1016/j.molcel.2008.07.020.

Herman, J.A., Miller, M.P. and Biggins, S. (2020) ‘chTOG is a conserved mitotic error correction factor’, eLife. Edited by S. Hauf et al., 9, p. e61773. Available at: 10.7554/eLife.61773.

Holder, J., Mohammed, S. and Barr, F.A. (2020) ‘Ordered dephosphorylation initiated by the selective proteolysis of cyclin B drives mitotic exit’, eLife, 9, p. e59885. Available at: 10.7554/eLife.59885.

Hsu, K.-S. and Toda, T. (2011) ‘Ndc80 internal loop interacts with Dis1/TOG to ensure proper kinetochore-spindle attachment in fission yeast’, Current biology: CB, 21(3), pp. 214–220. Available at: 10.1016/j.cub.2010.12.048.

Hu, C.-K., et al. (2012) ‘Plk1 negatively regulates PRC1 to prevent premature midzone formation before cytokinesis’, Molecular Biology of the Cell. Edited by Y.-L. Wang, 23(14), pp. 2702–2711. Available at: 10.1091/mbc.e12-01-0058.

Humphrey, L., Felzer-Kim, I. and Joglekar, A.P. (2018) ‘Stu2 acts as a microtubule destabilizer in metaphase budding yeast spindles’, Molecular Biology of the Cell, 29(3), pp. 247–255. Available at: 10.1091/mbc.E17-08-0494.

Játiva, S. et al. (2019) ‘Cdc14 activation requires coordinated Cdk1-dependent phosphorylation of Net1 and PP2A–Cdc55 at anaphase onset’, Cellular and Molecular Life Sciences, 76(18), pp. 3601– 3620. Available at: 10.1007/s00018-019-03086-5.

Jenni, S. and Harrison, S.C. (2018) ‘Structure of the DASH/Dam1 complex shows its role at the yeast kinetochore-microtubule interface’, Science, 360(6388), pp. 552–558. Available at: 10.1126/science.aar6436.

Joglekar, A.P., Bloom, K. and Salmon, E.D. (2009) ‘In vivo protein architecture of the eukaryotic kinetochore with nanometer scale accuracy’, Current biology: CB, 19(8), pp. 694–699. Available at: 10.1016/j.cub.2009.02.056.

Jonasson, E.M. et al. (2016) ‘Zds1/Zds2–PP2ACdc55 complex specifies signaling output from Rho1 GTPase’, Journal of Cell Biology, 212(1), pp. 51–61. Available at: 10.1083/jcb.201508119.

Khmelinskii, A. et al. (2007) ‘Cdc14-regulated midzone assembly controls anaphase B’, The Journal of Cell Biology, 177(6), pp. 981–993. Available at: 10.1083/jcb.200702145.

Kosco, K.A. et al. (2001) ‘Control of microtubule dynamics by Stu2p is essential for spindle orientation and metaphase chromosome alignment in yeast’, Molecular biology of the cell, 12(9), pp. 2870–2880.

Lanz, M.C. et al. (2021) ‘In-depth and 3-dimensional exploration of the budding yeast phosphoproteome’, EMBO reports, 22(2). Available at: 10.15252/embr.202051121.

Li, S., Garcia-Rodriguez, L.J. and Tanaka, T.U. (2023) ‘Chromosome biorientation requires Aurora B’s spatial separation from its outer kinetochore substrates, but not its turnover at kinetochores’, Current Biology, 33(21), pp. 4557–4569.e3. Available at: 10.1016/j.cub.2023.09.006.

Lim, W.M. et al. (2024) ‘Regulation of minimal spindle midzone organization by mitotic kinases’, Nature Communications, 15(1), p. 9213. Available at: 10.1038/s41467-024-53500-1.

Liu, H. and Naismith, J.H. (2008) ‘An efficient one-step site-directed deletion, insertion, single and multiple-site plasmid mutagenesis protocol’, BMC Biotechnology, 8(1), p. 91. Available at: 10.1186/1472-6750-8-91.

London, N. et al. (2012) ‘Phosphoregulation of Spc105 by Mps1 and PP1 Regulates Bub1 Localization to Kinetochores’, Current Biology, 22(10), pp. 900–906. Available at: 10.1016/j.cub.2012.03.052.

Longtine, M.S. et al. (1998) ‘Additional modules for versatile and economical PCR-based gene deletion and modification in Saccharomyces cerevisiae’, Yeast, 14(10), pp. 953–961. Available at: 10.1002/(SICI)1097-0061(199807)14:10<953::AID-YEA293>3.0.CO;2-U.

Ma, L. et al. (2007) ‘Spc24 and Stu2 promote spindle integrity when DNA replication is stalled’, Molecular Biology of the Cell, 18(8), pp. 2805–2816. Available at: 10.1091/mbc.E06-09-0882.

Marco, E. et al. (2013) ‘S. cerevisiae chromosomes biorient via gradual resolution of syntely between S phase and anaphase’, Cell, 154(5), pp. 1127–1139. Available at: 10.1016/j.cell.2013.08.008.

Miller, M.P. et al. (2019) ‘Kinetochore-associated Stu2 promotes chromosome biorientation in vivo’, PLOS Genetics, 15(10), p. e1008423. Available at: 10.1371/journal.pgen.1008423.

Miller, M.P., Asbury, C.L. and Biggins, S. (2016) ‘A TOG Protein Confers Tension Sensitivity to Kinetochore-Microtubule Attachments’, Cell, 165(6), pp. 1428–1439. Available at: 10.1016/j.cell.2016.04.030.

Moyano-Rodríguez, Y. et al. (2022) ‘PP2A-Cdc55 phosphatase regulates actomyosin ring contraction and septum formation during cytokinesis’, Cellular and Molecular Life Sciences, 79(3), p. 165. Available at: 10.1007/s00018-022-04209-1.

Moyano-Rodriguez, Y. and Queralt, E. (2019) ‘PP2A Functions during Mitosis and Cytokinesis in Yeasts’, International Journal of Molecular Sciences, 21(1), p. 264. Available at: 10.3390/ijms21010264.

Neef, R. et al. (2007) ‘Choice of Plk1 docking partners during mitosis and cytokinesis is controlled by the activation state of Cdk1’, Nature Cell Biology, 9(4), pp. 436–444. Available at: 10.1038/ncb1557.

Nishimura, K. et al. (2009) ‘An auxin-based degron system for the rapid depletion of proteins in nonplant cells’, Nature methods, 6(12), pp. 917–922. Available at: 10.1038/nmeth.1401.

Ólafsson, G. and Thorpe, P.H. (2015) ‘Synthetic physical interactions map kinetochore regulators and regions sensitive to constitutive Cdc14 localization’, Proceedings of the National Academy of Sciences, 112(33), pp. 10413–10418. Available at: 10.1073/pnas.1506101112.

Ólafsson, G. and Thorpe, P.H. (2020) ‘Polo kinase recruitment via the constitutive centromere-associated network at the kinetochore elevates centromeric RNA’, PLOS Genetics. Edited by B.A. Sullivan, 16(8), p. e1008990. Available at: 10.1371/journal.pgen.1008990.

Örd, M. et al. (2020) ‘Proline-Rich Motifs Control G2-CDK Target Phosphorylation and Priming an Anchoring Protein for Polo Kinase Localization’, Cell Reports, 31(11), p. 107757. Available at: 10.1016/j.celrep.2020.107757.

Park, C.J. et al. (2008) ‘Requirement for the Budding Yeast Polo Kinase Cdc5 in Proper Microtubule Growth and Dynamics’, Eukaryotic Cell, 7(3), pp. 444–453. Available at: 10.1128/EC.00283-07.

Pinsky, B.A. et al. (2006) ‘The Ipl1-Aurora protein kinase activates the spindle checkpoint by creating unattached kinetochores’, Nature Cell Biology, 8(1), pp. 78–83. Available at: 10.1038/ncb1341.

Queralt, E. et al. (2006) ‘Downregulation of PP2ACdc55 Phosphatase by Separase Initiates Mitotic Exit in Budding Yeast’, Cell, 125(4), pp. 719–732. Available at: 10.1016/j.cell.2006.03.038.

Santos, A., Wernersson, R. and Jensen, L.J. (2015) ‘Cyclebase 3.0: a multi-organism database on cell-cycle regulation and phenotypes’, Nucleic Acids Research, 43(D1), pp. D1140–D1144. Available at: 10.1093/nar/gku1092.

Sarangapani, K.K. et al. (2013) ‘Phosphoregulation promotes release of kinetochores from dynamic microtubules via multiple mechanisms’, Proceedings of the National Academy of Sciences of the United States of America, 110(18), pp. 7282–7287. Available at: 10.1073/pnas.1220700110.

Severin, F. et al. (2001) ‘Stu2 Promotes Mitotic Spindle Elongation in Anaphase’, The Journal of Cell Biology, 153(2), pp. 435–442. Available at: 10.1083/jcb.153.2.435.

Sharma, P. et al. (2019) ‘A cryptic hydrophobic pocket in the polo-box domain of the polo-like kinase PLK1 regulates substrate recognition and mitotic chromosome segregation’, Scientific Reports, 9(1), p. 15930. Available at: 10.1038/s41598-019-50702-2.

Singh, P. et al. (2021) ‘BUB1 and CENP-U, Primed by CDK1, Are the Main PLK1 Kinetochore Receptors in Mitosis’, Molecular Cell, 81(1), pp. 67–87.e9. Available at: 10.1016/j.molcel.2020.10.040.

Stangier, M.M. et al. (2018) ‘Structure-Function Relationship of the Bik1-Bim1 Complex’, Structure, 26(4), pp. 607–618.e4. Available at: 10.1016/j.str.2018.03.003.

Sundin, L.J.R., Guimaraes, G.J. and DeLuca, J.G. (2011) ‘The NDC80 complex proteins Nuf2 and Hec1 make distinct contributions to kinetochore–microtubule attachment in mitosis’, Molecular Biology of the Cell. Edited by D.G. Drubin, 22(6), pp. 759–768. Available at: 10.1091/mbc.e10-08-0671.

Tang, N.H. et al. (2013) ‘The internal loop of fission yeast Ndc80 binds Alp7/TACC-Alp14/TOG and ensures proper chromosome attachment’, Molecular Biology of the Cell, 24(8), pp. 1122–1133. Available at: 10.1091/mbc.E12-11-0817.

Touati, S.A. et al. (2019) ‘Cdc14 and PP2A Phosphatases Cooperate to Shape Phosphoproteome Dynamics during Mitotic Exit’, Cell Reports, 29(7), pp. 2105–2119.e4. Available at: 10.1016/j.celrep.2019.10.041.

Towbin, H., Staehelin, T. and Gordon, J. (1979) ‘Electrophoretic transfer of proteins from polyacrylamide gels to nitrocellulose sheets: procedure and some applications.’, Proceedings of the National Academy of Sciences, 76(9), pp. 4350–4354. Available at: 10.1073/pnas.76.9.4350.

Tseng, W.-C. et al. (2008) ‘A novel megaprimed and ligase-free, PCR-based, site-directed mutagenesis method’, Analytical Biochemistry, 375(2), pp. 376–378. Available at: 10.1016/j.ab.2007.12.013.

Ubersax, J.A. et al. (2003) ‘Targets of the cyclin-dependent kinase Cdk1’, Nature, 425(6960), pp. 859–864. Available at: 10.1038/nature02062.

Uhlmann, F. et al. (2000) ‘Cleavage of Cohesin by the CD Clan Protease Separin Triggers Anaphase in Yeast’, Cell, 103(3), pp. 375–386. Available at: 10.1016/S0092-8674(00)00130-6.

Usui, T. et al. (2003) ‘The XMAP215 homologue Stu2 at yeast spindle pole bodies regulates microtubule dynamics and anchorage’, The EMBO journal, 22(18), pp. 4779–4793. Available at: 10.1093/emboj/cdg459.

Valverde, R., Ingram, J. and Harrison, S.C. (2016) ‘Conserved Tetramer Junction in the Kinetochore Ndc80 Complex’, Cell Reports, 17(8), pp. 1915–1922. Available at: 10.1016/j.celrep.2016.10.065.

Van Der Vaart, B. et al. (2017) ‘TORC1 signaling exerts spatial control over microtubule dynamics by promoting nuclear export of Stu2’, Journal of Cell Biology, 216(11), pp. 3471–3484. Available at: 10.1083/jcb.201606080.

Vasileva, V. et al. (2017) ‘Molecular mechanisms facilitating the initial kinetochore encounter with spindle microtubules’, The Journal of Cell Biology, 216(6), pp. 1609–1622. Available at: 10.1083/jcb.201608122.

Wang, C. et al. (2020) ‘GPS 5.0: An Update on the Prediction of Kinase-specific Phosphorylation Sites in Proteins’, Genomics, Proteomics & Bioinformatics, 18(1), pp. 72–80. Available at: 10.1016/j.gpb.2020.01.001.

Wang, P.J. and Huffaker, T.C. (1997) ‘Stu2p: A microtubule-binding protein that is an essential component of the yeast spindle pole body’, The Journal of Cell Biology, 139(5), pp. 1271–1280. Available at: 10.1083/jcb.139.5.1271.

Yaakov, G., Thorn, K. and Morgan, D.O. (2012) ‘Separase Biosensor Reveals that Cohesin Cleavage Timing Depends on Phosphatase PP2ACdc55 Regulation’, Developmental Cell, 23(1), pp. 124–136. Available at: 10.1016/j.devcel.2012.06.007.

Yellman, C.M. and Burke, D.J. (2006) ‘The Role of Cdc55 in the Spindle Checkpoint Is through Regulation of Mitotic Exit in *Saccharomyces cerevisiae*’, Molecular Biology of the Cell, 17(2), pp. 658–666. Available at: 10.1091/mbc.e05-04-0336.

Zahm, J.A., et al. (2021) ‘Structural basis of Stu2 recruitment to yeast kinetochores’, eLife. Edited by J.G. DeLuca, 10, p. e65389. Available at: 10.7554/eLife.65389.

Zaytsev, A.V., et al. (2015) ‘Multisite phosphorylation of the NDC80 complex gradually tunes its microtubule-binding affinity’, Molecular Biology of the Cell. Edited by T. Surrey, 26(10), pp. 1829–1844. Available at: 10.1091/mbc.E14-11-1539.

Zhu, J. et al. (2015) ‘Single-Cell Based Quantitative Assay of Chromosome Transmission Fidelity’, G3 Genes|Genomes|Genetics, 5 (6), pp. 1043–1056. Available at: 10.1534/g3.115.017913.

Zimniak, T. et al. (2012) ‘Spatiotemporal Regulation of Ipl1/Aurora Activity by Direct Cdk1 Phosphorylation’, Current Biology, 22(9), pp. 787–793. Available at: 10.1016/j.cub.2012.03.007.

